# A cerebrospinal fluid proteomic clock reveals opposing brain-aging programs and predicts neurological disease progression

**DOI:** 10.64898/2026.06.26.734784

**Authors:** Shenrui Xu, Yanying Guo, Ke Fang, Suli Li, Tingting Wang, Yang li, Mengyi Zhang, Hongyan Li, Tao Xu, Zhichao Miao, Yining Yang, Zonghong Li

## Abstract

Aging is the risk factor for neurological disease, yet the molecular architecture of human brain aging remains poorly defined. Here, we analyzed more than 10,000 cerebrospinal fluid (CSF) proteomes across multiple cohorts and proteomic platforms to developed a 249-protein CSF aging clock that accurately predicts age and generalizes across independent datasets. CSF brain-age acceleration was increased across diverse neurological diseases, associated with blood-brain barrier (BBB) dysfunction, and predictive of longitudinal cognitive decline, neuroimaging progression, and dementia conversion. A simplified 30-protein panel retained similar prognostic performance. Biologically, clock proteins resolved two opposing programs: pro-aging activation of immune, vascular/BBB, ECM and coagulation pathways marked by CHI3L1, CD14, VWF, LRG1 and LTBP2, and anti-aging collapse of neuronal, synaptic, axonal and structural-maintenance programs marked by NPTX2, COL1A2, NID1, CDH8 and PENK. Brain-wide single-cell and regional mapping linked these programs to disease-vulnerable compartments. These findings establish a molecular framework for biological brain aging and neurological disease vulnerability.

## Introduction

Aging is the strongest risk factor for nearly all major neurological disorders, including Alzheimer’s disease (AD), Parkinson’s disease (PD), multiple sclerosis (MS), stroke, and neurodegenerative dementias^1,2^. Despite their distinct etiologies, these disorders share common age-associated pathological features, suggesting that biological brain aging may represent a convergent mechanism underlying neurological vulnerability^3,4^. However, the molecular programs that define human brain aging remain poorly understood.

Current approaches for measuring biological aging primarily rely on DNA methylation, transcriptomic signatures, or circulating plasma proteins^5–8^. Although these biomarkers have substantially advanced aging research, they incompletely capture the molecular processes occurring within the central nervous system^9^. CSF, which directly interfaces with the brain parenchyma and reflects ongoing neurobiological processes, provides a unique opportunity to characterize molecular aging within the human brain^10^. Early CSF proteomic studies identified age-associated alterations in inflammatory and injury-response proteins, while more recent large-scale analyses revealed widespread effects of age, sex, APOE ε4 genotype, and amyloid pathology on the CSF proteome^8,11^. Collectively, these studies demonstrate that aging exerts profound effects on the molecular composition of CSF. However, whether these age-related proteomic alterations can be integrated into a quantitative measure of biological brain aging and, more importantly, whether they reveal conserved molecular programs underlying human brain aging remain unknown.

Here, we address this gap by developing a CSF proteomic aging clock from a large discovery cohort and validating it across more than 10,000 individuals, three orthogonal proteomic platforms, and multiple neurological conditions. We show that CSF brain-age acceleration is elevated across diverse neurological diseases, associates with BBB dysfunction, and predicts future neurodegenerative progression. Mechanistically, the aging clock resolves two opposing molecular programs characterized by activation of inflammatory-neurovascular remodeling pathways and decline of neuronal-maintenance networks. Integration with a human brain single-cell atlas localized region- and cell-type-specific clock further localizes these programs to disease-vulnerable neurovascular and neuronal circuits and reveals a non-linear trajectory of brain aging characterized by a prominent midlife remodeling wave. Together, this work establishes a scalable, cross-platform CSF proteomic framework for quantifying central nervous system (CNS) biological aging and resolving disease-specific aging trajectories across the neurological disease spectrum.

## Results

### Machine learning of CSF proteomic across multiple cohorts and proteomic platforms identifies a 249-protein aging clock

An overview of the study design is presented in Fig.1. To establish a CSF-specific measure of biological brain aging, we analyzed more than 10,000 CSF proteomes generated across multiple cohorts and proteomic platforms. We first developed a CSF proteomic aging clock in neurologically normal individuals and subsequently evaluated its robustness across independent cohorts, orthogonal proteomic technologies, and diverse neurological diseases. In parallel, we integrated clock proteins with a human brain single-cell atlas comprising 2.25 million cells from 1,632 healthy individuals across thirteen brain regions to investigate their cellular and anatomical context.

**Figure. 1.**
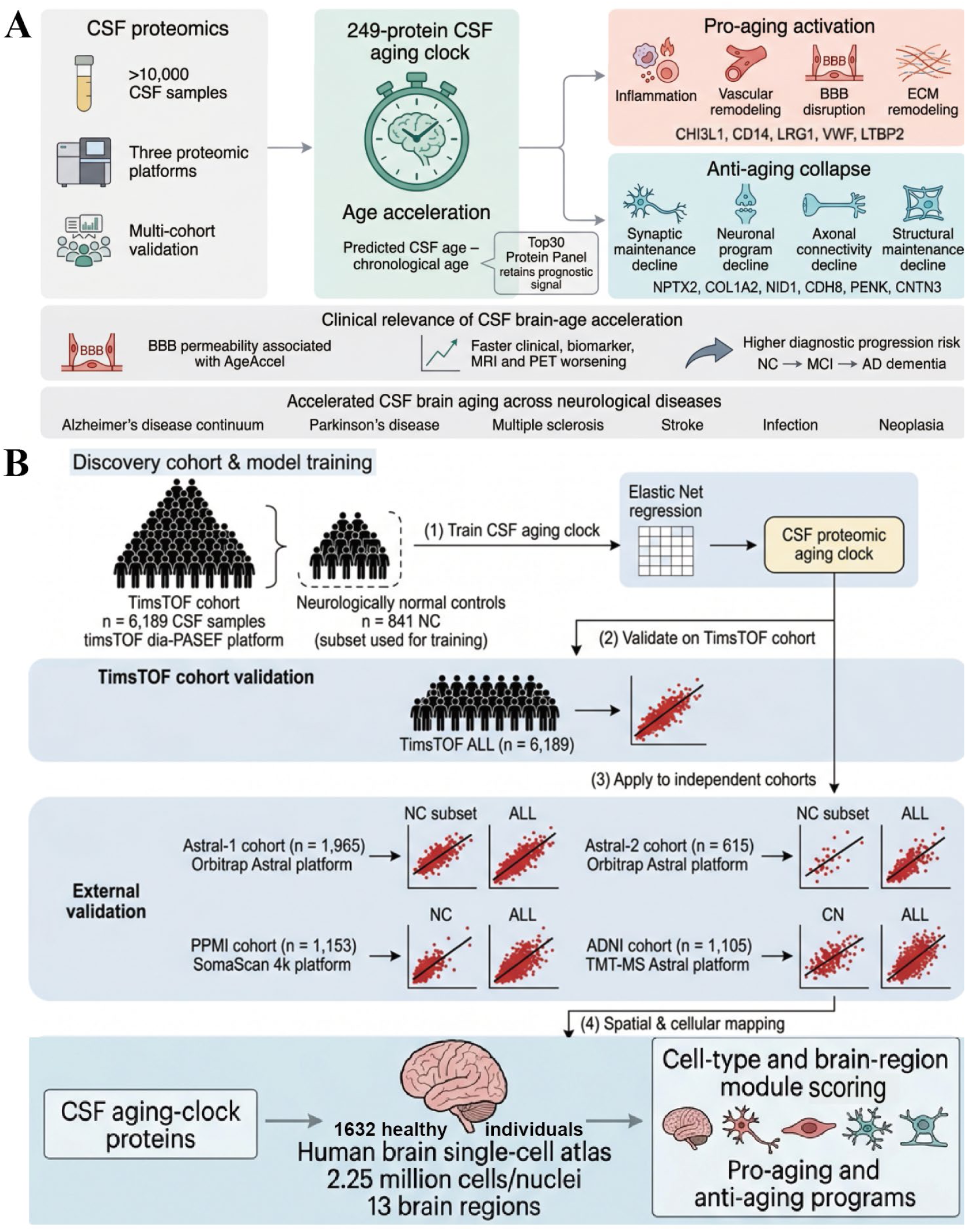
Study overview and analytical framework for defining biological brain aging using CSF proteomics. **(A)** Conceptual overview of the study. A CSF proteomic aging clock was developed to quantify biological brain age and derive CSF brain-age acceleration (AgeAccel), defined as predicted CSF age minus chronological age. The framework was used to characterize biological brain aging across diverse neurological disorders, including the Alzheimer’s disease continuum, Parkinson’s disease, multiple sclerosis, stroke, infection, and neoplasia. Downstream analyses investigated the molecular architecture of brain aging, its relationship to blood–brain barrier dysfunction and disease progression, its anatomical and cellular origins within the human brain, and its temporal dynamics across the lifespan. **(B)** Cohort design and validation workflow. The CSF aging clock was trained using neurologically normal controls (n = 841) from the timsTOF discovery cohort (n = 6,189) profiled by dia-PASEF mass spectrometry. Model performance was evaluated in the full timsTOF cohort and independently validated across four external cohorts measured using orthogonal proteomic platforms, including Orbitrap Astral (Astral-1, n = 1,965; Astral-2, n = 615), SomaScan 4k aptamer proteomics (PPMI, n = 1,153), and TMT-MS Orbitrap Astral proteomics (ADNI, n = 1,105). The resulting clock was subsequently used to assess disease-associated age acceleration, longitudinal disease progression, molecular aging programs, genetic enrichment, and regional and cellular localization using a human brain single-cell atlas comprising 2.25 million cells from 1,632 healthy individuals across 13 brain regions.

Using 841 neurologically normal controls from the timsTOF cohort (n = 6,189)^12^, elastic-net regression identified a 249-protein CSF aging signature (Supplementary Table 1). The resulting model accurately predicted chronological age in the training population (r = 0.937, MAE = 3.82 years; Fig. 2A). Importantly, age-predictive performance remained robust in independent cohorts profiled using the Orbitrap Astral platform, achieving correlations of r = 0.900 (MAE = 5.8 years) in Astral-1 and r = 0.931 (MAE = 4.4 years) in Astral-2 (Fig. 2B, C). These findings demonstrate that the identified proteomic aging signature generalizes beyond the training platform and captures reproducible age-associated biological variation.

**Figure 2.**
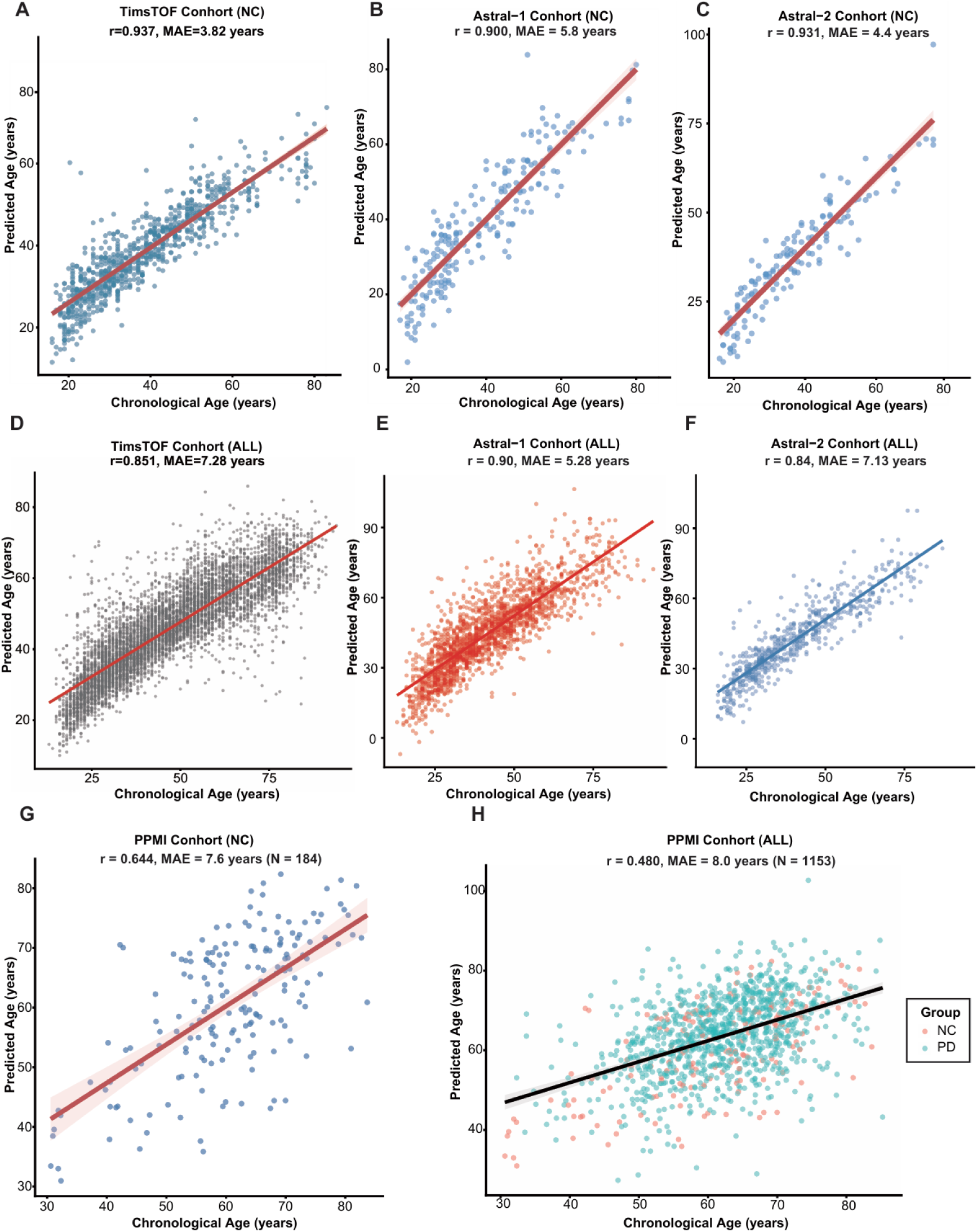
Development and cross-platform validation of the CSF proteomic aging clock. An elastic-net regression model was trained using CSF proteomic profiles from neurologically normal controls in the timsTOF discovery cohort, yielding a 249-protein CSF aging clock. Scatter plots show predicted age versus chronological age. Solid lines indicate linear regression fits. Pearson correlation coefficients (*r*) and mean absolute errors (MAE) are shown in each panel. **(A–C)** Age-prediction performance in neurologically normal controls from the timsTOF discovery cohort (**A**, *n* = 841), Astral-1 cohort (**B**, *n* = 221), and Astral-2 cohort (**C**, *n* = 155). **(D–F)** Validation in all available samples from the timsTOF cohort (**D**, *n* = 6,189), Astral-1 cohort (**E**, *n* = 1,965), and Astral-2 cohort (**F**, *n* = 615), including both neurologically normal controls and neurological disease cases. **(G, H)** External validation in the Parkinson’s Progression Markers Initiative (PPMI) cohort profiled using SomaScan aptamer-based proteomics. Model performance is shown in healthy controls (**G**, *n* = 184) and the full PPMI cohort (**H**, *n* = 1,153). Points in panel H are colored according to diagnostic group.

When applied to all available samples, including neurological disease populations, predicted CSF age remained strongly associated with chronological age in the full timsTOF cohort (r = 0.851, MAE = 7.28 years), Astral-1 cohort (r = 0.900, MAE = 5.28 years), and Astral-2 cohort (r = 0.840, MAE = 7.13 years) (Fig. 2D–F). Notably, prediction error increased in disease-enriched cohorts relative to neurologically normal controls, suggesting that deviations from the normative aging trajectory may reflect biologically meaningful alterations associated with neurological pathology rather than technical variation alone.

To further evaluate platform transferability, we applied the model to the Parkinson’s Progression Markers Initiative (PPMI) cohort profiled using the SomaScan aptamer-based proteomics platform. Following cohort-specific calibration, predicted CSF age remained significantly associated with chronological age in healthy controls (r = 0.644, MAE = 7.6 years; n = 184; Fig. 2G). Across the full PPMI cohort, including Parkinson’s disease cases, the association remained significant albeit attenuated (r = 0.480, MAE = 8.0 years; n = 1,153; Fig. 2H). Similarly, in the independent ADNI cohort profiled using TMT-based mass spectrometry, predicted age remained positively associated with chronological age across cognitively normal controls, mild cognitive impairment, and Alzheimer’s disease dementia (r = 0.407, MAE = 6.75 years; n = 1,105; Supplementary Fig. 1). Despite substantial differences in disease composition, cohort structure, and proteomic technology, the clock consistently captured age-related proteomic architecture across all validation datasets.

Sex-stratified analyses further supported the robustness of the model. Among neurologically normal controls, age prediction accuracy was comparable between females (r = 0.933, MAE = 3.61 years; n = 620) and males (r = 0.940, MAE = 4.41 years; n = 221; Supplementary Fig. 2A). Similar performance was observed in the full timsTOF cohort, including disease samples, for both females (r = 0.851, MAE = 6.85 years; n = 3,442) and males (r = 0.845, MAE = 7.81 years; n = 2,747; Supplementary Fig. 2B). These findings indicate that the clock is not driven by sex-specific prediction bias and remains stable across demographic strata.

Collectively, these results establish a 249-protein CSF aging clock that captures a conserved molecular architecture of brain aging reproducibly across independent cohorts, neurological diseases, and orthogonal proteomic platforms.

### CSF proteomic aging clock reveals biological brain age acceleration in diverse neurological diseases

We next investigated whether deviation from the normative CSF aging trajectory was associated with neurological disease. Using age-adjusted CSF age acceleration as a measure of biological brain aging, we observed widespread acceleration across diverse neurological disorders in the timsTOF cohort (Fig. 3A). Relative to neurologically normal controls, individuals with neurodegenerative disease, multiple sclerosis, other autoimmune neurological disorders, neoplasia, infectious disease, stroke, and other neurological conditions exhibited elevated CSF age acceleration. Although substantial inter-individual heterogeneity was present within diagnostic groups, the overall distribution of age acceleration was consistently shifted toward older biological brain age across most disease categories.

**Figure 3.**
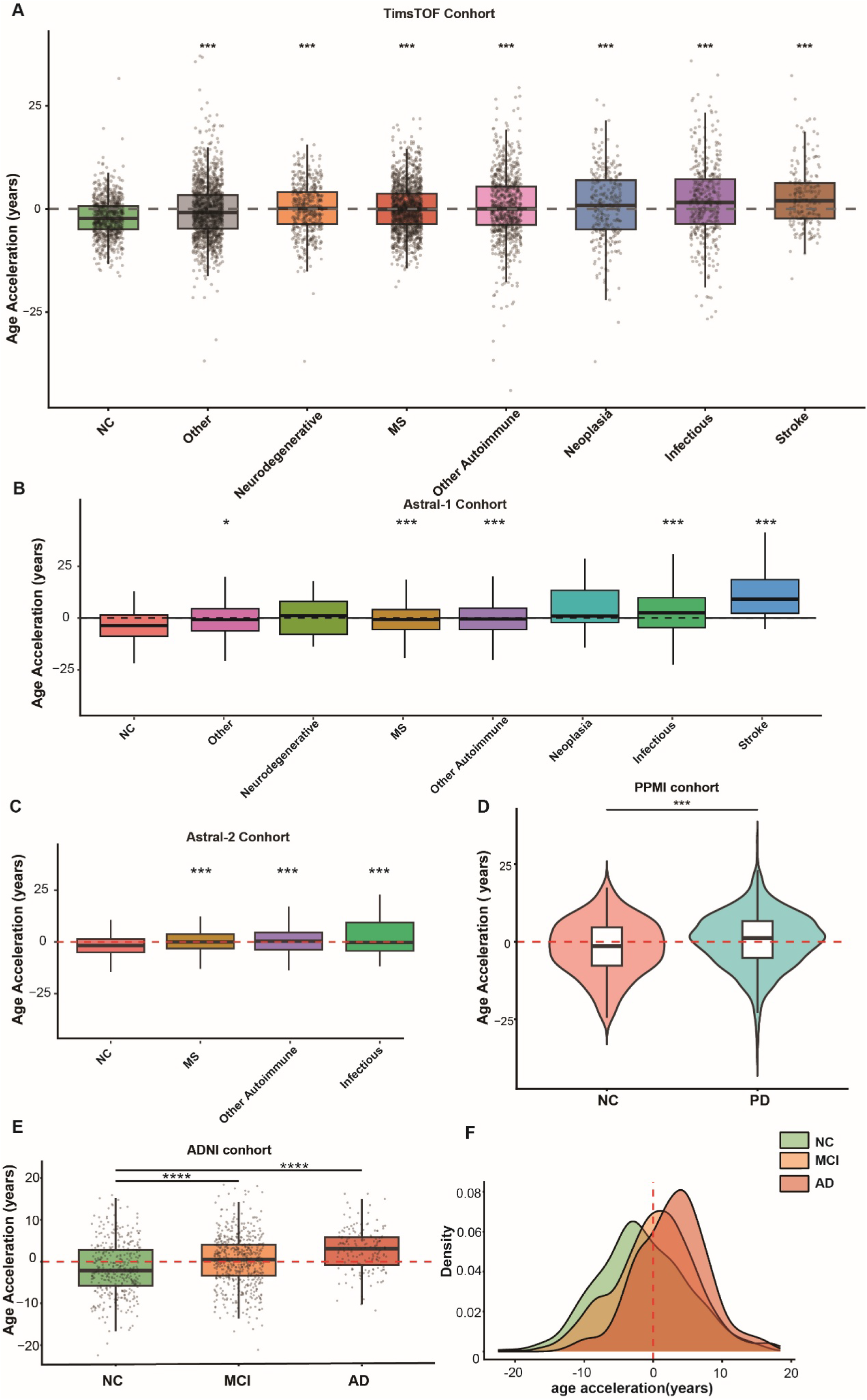
CSF brain-age acceleration across neurological diseases, cohorts, and proteomic platforms. **(A)** Brain-age acceleration (AgeAccel) across diagnostic groups in the timsTOF cohort (*n* = 6,189), including neurologically normal controls (NC), multiple sclerosis (MS), other CNS diseases, other autoimmune neurological diseases, neurodegenerative diseases, infectious diseases, neoplasia, and ischemic stroke. **(B)** AgeAccel in the Astral-1 validation cohort (*n* = 1,965), comprising a stratified subset of samples re-profiled using the Orbitrap Astral platform. **(C)** AgeAccel in the independent Astral-2 cohort (*n* = 615), including MS, autoimmune neurological diseases, infectious diseases, and controls. **(D)** AgeAccel in the PPMI cohort (*n* = 1,153), comparing healthy controls (HC, *n* = 184) and Parkinson’s disease (PD, *n* = 969) participants profiled using SomaScan proteomics. **(E)** AgeAccel across the Alzheimer’s disease continuum in the ADNI cohort (*n* = 1,105), including cognitively normal controls (NC, *n* = 379), mild cognitive impairment (MCI, *n* = 562), and Alzheimer’s disease dementia (AD, *n* = 164). **(F)** Density distributions of AgeAccel across NC, MCI, and AD groups in ADNI. Box plots show the median (center line), interquartile range (box), and 1.5× IQR (whiskers). Statistical significance was assessed using two-sided Wilcoxon rank-sum tests with Benjamini-Hochberg false-discovery-rate correction. *P* values are denoted as *p < 0.05, **p < 0.01, ***p < 0.001, ****p < 0.0001; ns, not significant.

These findings were independently replicated in the Orbitrap Astral validation cohorts. In Astral-1, multiple sclerosis, other autoimmune neurological diseases, infectious diseases, and stroke displayed significantly elevated age acceleration relative to controls (Fig. 3B). Similar results were observed in Astral-2, where all available disease groups, including multiple sclerosis, autoimmune neurological disease, and infectious disease, showed positive shifts in biological age acceleration compared with neurologically normal individuals (Fig. 3C). The reproducibility of these findings across independent cohorts and mass-spectrometry platforms indicates that disease-associated CSF aging acceleration reflects a robust biological signal rather than a platform-specific phenomenon.

External validation cohorts further extended these observations to major neurodegenerative diseases. In the PPMI cohort, individuals with Parkinson’s disease exhibited significantly greater CSF age acceleration than healthy controls (Fig. 3D), indicating accelerated biological brain aging in synucleinopathy. In the ADNI cohort, age acceleration increased progressively along the Alzheimer’s disease continuum, with cognitively normal controls showing the lowest values, individuals with mild cognitive impairment displaying intermediate acceleration, and patients with Alzheimer’s disease exhibiting the greatest deviation from the normative aging trajectory (Fig. 3E). Density distributions demonstrated a corresponding rightward shift from cognitively normal controls to mild cognitive impairment and Alzheimer’s disease (Fig. 3F), consistent with a gradual increase in biological brain aging across disease stages. Together, these findings suggest that CSF age acceleration captures both generalized neurological disease burden and progressive neurodegenerative deterioration.

Sex-stratified analyses further supported the robustness of the disease association while revealing quantitative differences between females and males. Among females, all evaluated disease categories demonstrated significantly elevated age acceleration relative to sex-matched controls (Supplementary Fig. 3A). In males, significant acceleration was observed in multiple sclerosis, autoimmune neurological disease, infectious disease, and stroke, whereas other neurological disorders, neurodegenerative diseases, and neoplasia did not reach statistical significance in the available comparisons (Supplementary Fig. 3B). These results indicate that accelerated CSF brain aging is evident in both sexes, although its magnitude and disease specificity may vary according to biological sex. Collectively, these findings demonstrate that CSF age acceleration represents a reproducible marker of accelerated biological brain aging that is consistently elevated across a broad spectrum of neurological diseases and progressively increases along major neurodegenerative disease trajectories.

### CSF brain-age acceleration is associated with BBB dysfunction and predicts longitudinal disease progression

We next investigated whether CSF brain-age acceleration captured clinically relevant features of CNS vulnerability. In the discovery clinical cohort, individuals with elevated BBB permeability, defined using age-adjusted cerebrospinal fluid/serum albumin quotient (QAlb) thresholds, exhibited significantly greater CSF brain-age acceleration (AgeAccel) than those with normal BBB function (difference = 3.26 years; n = 6,139; P < 0.0001; Fig. 4A). Consistent with a dose-dependent relationship, AgeAccel increased progressively across QAlb tertiles, with individuals in the highest tertile showing substantially greater acceleration than those in the lowest tertile (difference = 4.50 years; P < 0.0001; Fig. 4B). Similar associations were observed for multiple BBB- and intrathecal immunity-related measures, including QAlb, cerebrospinal fluid/serum IgG quotient (QIgG), CSF albumin, CSF IgG, and oligoclonal bands, all of which demonstrated significant positive age- and sex-adjusted correlations with AgeAccel (FDR < 0.001; Supplementary Fig. 4). Together, these findings indicate that accelerated CSF brain aging is closely linked to BBB dysfunction and neuroinflammatory barrier pathology.

**Figure 4.**
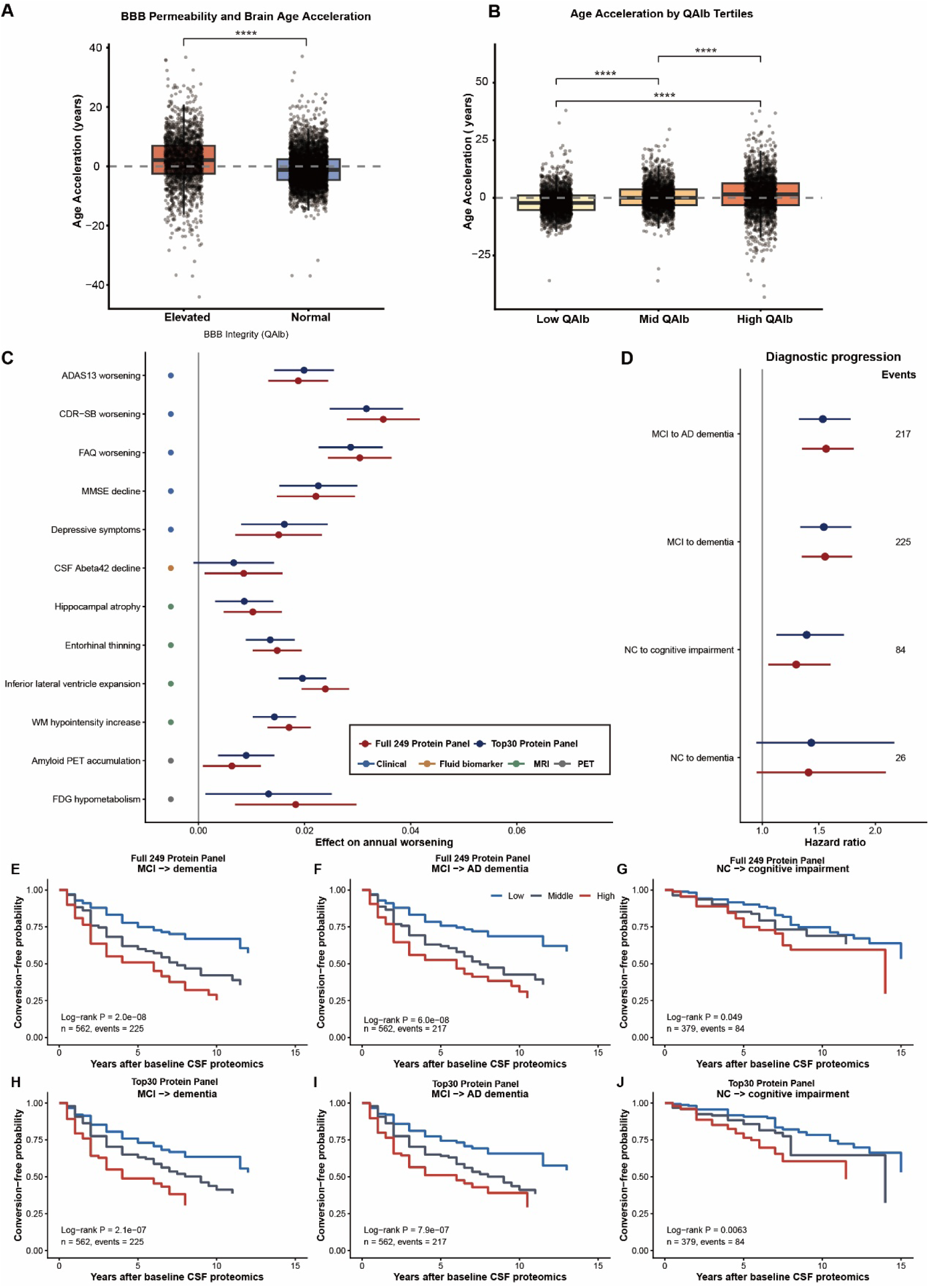
CSF AgeAccel is associated with BBB dysfunction and longitudinal disease progression. **(A)** AgeAccel in individuals with normal and elevated BBB permeability, defined using age-adjusted QAlb thresholds. **(B)** AgeAccel across QAlb tertiles. Box plots show the median (center line), interquartile range (box), and 1.5× interquartile range (whiskers). Individual points represent CSF samples. Statistical significance was assessed using two-sided Wilcoxon rank-sum tests. ****P < 0.0001. **(C)** Associations between baseline AgeAccel and longitudinal change in clinical, fluid biomarker, MRI, and PET outcomes in ADNI. Points indicate the estimated interaction effect between baseline AgeAccel and follow-up time per 1-s.d. increase in AgeAccel. Horizontal bars indicate 95% confidence intervals. Positive effect sizes correspond to faster worsening after outcome orientation. Estimates are shown for both the full 249-protein clock and the reduced Top30 panel. **(D)** Cox proportional-hazards models for diagnostic progression in ADNI. Hazard ratios are shown per 1-s.d. increase in baseline AgeAccel after adjustment for age, sex, and education. Horizontal bars indicate 95% confidence intervals. Event numbers are shown on the right. **(E–G)** Kaplan–Meier analyses of conversion-free survival stratified by tertiles of baseline AgeAccel derived from the full 249-protein clock. **(H–J)** Kaplan–Meier analyses of conversion-free survival stratified by tertiles of baseline AgeAccel derived from the reduced Top30 panel. P values for survival analyses were calculated using two-sided log-rank tests.

We next examined whether baseline AgeAccel predicted future disease progression in the independent ADNI cohort. Higher baseline AgeAccel was associated with accelerated longitudinal deterioration across multiple clinical, molecular, neuroimaging, and functional outcomes (Fig. 4C). The strongest effects were observed for cognitive and functional decline, including worsening Clinical Dementia Rating–Sum of Boxes (CDR-SB), Functional Activities Questionnaire (FAQ), Mini-Mental State Examination (MMSE), and 13-item Alzheimer’s Disease Assessment Scale–Cognitive Subscale (ADAS13) scores. Elevated AgeAccel was also associated with more rapid hippocampal atrophy, entorhinal cortical thinning, inferior lateral ventricular enlargement, increased white-matter abnormalities, greater amyloid accumulation on PET imaging, and progressive cerebral hypometabolism measured by FDG-PET. These results suggest that accelerated biological brain aging is associated with coordinated deterioration across multiple dimensions of neurodegenerative disease.

Consistent with these observations, Cox proportional-hazards models adjusted for age, sex, and education demonstrated that higher baseline AgeAccel was a strong predictor of future diagnostic progression (Fig. 4D). For each one-standard-deviation increase in AgeAccel, the risk of progression from mild cognitive impairment (MCI) to Alzheimer’s disease dementia increased by 56% (HR = 1.56, 95% CI 1.35–1.81, P = 1.8 × 10⁻⁹), while the risk of progression from MCI to all-cause dementia increased by a similar magnitude (HR = 1.56, 95% CI 1.35–1.79, P = 1.3 × 10⁻⁹). Higher AgeAccel also predicted conversion from cognitively normal status to cognitive impairment (HR = 1.30, 95% CI 1.05–1.60, P = 0.015). Kaplan–Meier analyses further demonstrated progressive stratification of conversion-free survival across AgeAccel tertiles, with individuals in the highest tertile exhibiting the greatest risk of clinical progression for MCI-to-dementia, MCI-to-Alzheimer’s disease dementia, and cognitively normal-to-cognitive impairment transitions (log-rank P = 2.0 × 10⁻⁸, 6.0 × 10⁻⁸, and 0.049, respectively; Fig. 4E–G).

To evaluate translational feasibility, we derived a simplified 30-protein panel from the full 249-protein aging clock. Despite substantial reduction in model complexity, the Top30 Protein Panel preserved the longitudinal associations observed for the full model across clinical, biomarker, and neuroimaging outcomes (Fig. 4C). Baseline Top30 AgeAccel remained strongly associated with diagnostic progression, predicting conversion from MCI to Alzheimer’s disease dementia (HR = 1.54, 95% CI 1.32–1.78, P = 1.5 × 10⁻⁸), MCI to all-cause dementia (HR = 1.54, 95% CI 1.34–1.79, P = 4.9 × 10⁻⁹), and cognitively normal status to cognitive impairment (HR = 1.39, 95% CI 1.13–1.72, P = 0.0023; Fig. 4D). Kaplan–Meier analyses confirmed robust risk stratification across Top30 AgeAccel tertiles for all progression outcomes examined (Fig. 4H–J).

Collectively, these findings demonstrate that CSF brain-age acceleration is closely linked to BBB dysfunction, predicts future neurodegenerative progression across multiple biological and clinical domains, and can be captured using a clinically tractable 30-protein panel, highlighting its potential utility as a biomarker of biological brain aging and disease vulnerability.

### Clock proteins separated into coordinated pro-aging and anti-aging molecular programs

We next examined the biological architecture underlying the CSF proteomic aging clock. At the individual-protein level, clock proteins separated into two opposing axes when ranked against calibrated CSF brain-age acceleration (Fig. 5A). Proteins with positive clock coefficients formed a pro-aging activation axis, whereas proteins with negative coefficients formed an anti-aging collapse axis. This organization was not only a coefficient-level feature of the elastic-net model: protein-AgeAccel correlations were directionally concordant with model coefficients for 81.5% of clock proteins (Fig. 5B). Representative pro-aging proteins included CHI3L1, CD14, LRG1, VWF and LTBP2, linking higher CSF age acceleration to inflammatory, vascular and matrix-remodeling biology. Conversely, representative anti-aging proteins included NPTX2, NPTXR, CDH8, CDH10 and PTPRR, indicating loss of proteins associated with synaptic, neuronal and axonal maintenance.

**Figure 5.**
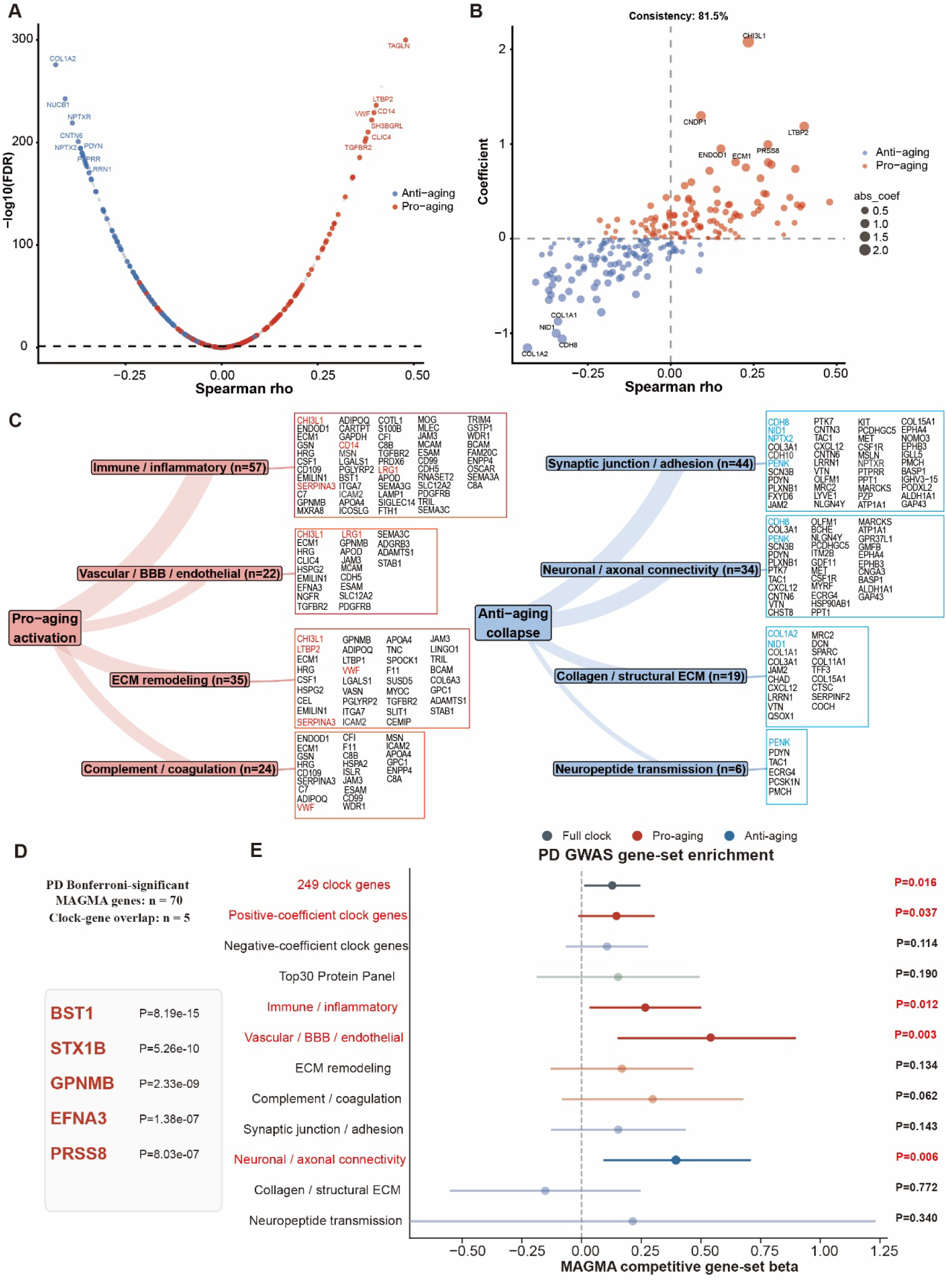
Dual-axis molecular architecture of the CSF proteomic aging clock and its convergence with Parkinson’s disease genetic susceptibility. **(A)** Associations between individual CSF aging-clock proteins and calibrated AgeAccel. The x axis shows Spearman correlation coefficients and the y axis shows −log10(FDR). Proteins with positive elastic-net coefficients are shown in red, whereas proteins with negative coefficients are shown in blue. Labelled proteins indicate representative members of the major clock-associated pathways. **(B)** Concordance between protein–AgeAccel correlations and elastic-net clock coefficients. Each point represents one clock protein; color indicates coefficient direction and point size corresponds to the absolute coefficient magnitude. The dashed vertical line indicates zero correlation. Overall directional concordance between model coefficients and AgeAccel-associated protein changes was 81.5%. Labelled proteins denote representative pro-aging proteins (CHI3L1, CD14, LRG1, VWF, and LTBP2) and anti-aging proteins (NPTX2, NPTXR, CDH8, CDH10, PTPRR, and COL1A1). **(C)** Functional organization of clock proteins into pro-aging and anti-aging molecular modules based on pathway-enrichment analyses. Pro-aging proteins were grouped into immune/inflammatory, vascular/BBB/endothelial, extracellular matrix (ECM)-remodeling, and complement/coagulation modules. Anti-aging proteins were grouped into synaptic junction/adhesion, neuronal/axonal-connectivity, collagen/structural ECM, and neuropeptide-transmission modules. Boxed protein lists indicate proteins assigned to each functional category; highlighted proteins denote representative module members. **(D)** Direct overlap between CSF aging-clock genes and Bonferroni-significant MAGMA genes identified in the Parkinson’s disease GWAS. Five clock genes overlapped with PD-associated susceptibility genes, including BST1, STX1B, GPNMB, EFNA3, and PRSS8. **(E)** Competitive MAGMA gene-set enrichment analyses of CSF aging-clock gene sets in the Parkinson’s disease GWAS. Points indicate MAGMA beta estimates and horizontal bars indicate 95% confidence intervals. Red labels denote nominally significant enrichments (P < 0.05). Gene sets include the full clock, pro-aging and anti-aging gene sets, and functional clock modules.

Functional grouping of clock proteins reinforced this two-axis structure. Pro-aging proteins were concentrated in immune/inflammatory, vascular/BBB/endothelial, extracellular matrix (ECM)-remodeling and complement/coagulation modules (Fig. 5C). These modules contained vascular and barrier-related proteins such as VWF, LTBP2, PDGFRB and CDH5, inflammatory or innate immune proteins such as CHI3L1, CD14, CSF1 and SERPINA3, and ECM-associated proteins including ECM1, HSPG2, ADAMTS1 and EMILIN1. In contrast, anti-aging proteins clustered into synaptic junction/adhesion, neuronal/axonal connectivity, collagen/structural ECM and neuropeptide-transmission modules. These modules included synaptic and neuronal proteins such as NPTX2, NPTXR, PTPRR, SCN3B, CNTN3, CNTN6, LRRN1 and PENK, together with structural proteins including COL1A1, COL1A2, NID1, DCN and SPARC. Thus, the clock did not simply track a generic inflammatory state; it captured the balance between activation of barrier-vascular-matrix programs and erosion of neuronal-maintenance programs. We further asked whether these pathway-defined proteins were dysregulated across neurological disease groups. Differential abundance analysis relative to neurologically normal controls showed a broad disease-associated shift in the same two directions (Supplementary Fig. 5A). Proteins belonging to the pro-aging modules, including SERPINA3, CD14, LRG1, VWF, ICAM2, SPOCK1, GPC1, MLEC and MCAM, tended to be increased across disease groups, with particularly prominent changes in stroke, neoplasia and infectious disease samples. By contrast, proteins belonging to the anti-aging modules, including NPTX2, NPTXR, CDH8, CDH10, PTPRR, CNTN3, CNTN6, EPHA4 and PENK, tended to be reduced across disease groups. The proteins highlighted in the heatmap overlapped with the pathway-defined proteins in Fig. 5C, indicating that disease-associated CSF proteomic changes recapitulated the same pro-aging and anti-aging programs identified from the clock architecture.

To test whether CSF aging-clock proteins intersect inherited susceptibility to neurodegenerative disease, we performed MAGMA gene-level and competitive gene-set enrichment analyses using PD and AD GWAS summary statistics. In the PD GWAS, direct overlap with Bonferroni-significant MAGMA genes identified five clock genes: BST1, STX1B, GPNMB, EFNA3 and PRSS8. All five belonged to the positive-coefficient pro-aging axis (Fig. 5D). At the gene-set level, the PD GWAS showed a broader nominal enrichment pattern across CSF aging-clock modules. The full 249-gene clock showed nominal enrichment (beta = 0.129, P = 0.016), as did positive-coefficient clock genes (beta = 0.146, P = 0.037). Among functional modules, immune/inflammatory genes (beta = 0.267, P = 0.0125), vascular/BBB/endothelial genes (beta = 0.525, P = 0.00296) and neuronal/axonal connectivity genes (beta = 0.401, P = 0.00554) showed positive enrichment signals (Fig. 5E). By contrast, in the AD GWAS, direct overlap was limited to two clock genes, BCAM and SERPINF2, and none of the clock-derived gene sets showed nominal enrichment (Supplementary Fig. 5B, C). These findings suggest that CSF aging-clock biology converges more strongly with PD than AD genetic susceptibility, particularly through pro-aging immune/vascular programs and neuronal/axonal connectivity pathways.

SHAP analyses further demonstrated that individual diseases achieved similar AgeAccel phenotypes through distinct combinations of clock proteins. For example, CHI3L1 and PRSS8 (PD susceptibility gene) contributed prominently to age acceleration in multiple sclerosis and neurodegenerative disorders, whereas extracellular matrix and structural proteins (COL1A2) contributed more strongly in neoplasia (Supplementary Fig. 6A-F). These observations suggest that diverse neurological disorders converge onto a common biological aging state through partially distinct molecular trajectories.

To determine whether the CSF clock captures a brain-specific aging program, we compared clock proteins with established blood-derived aging clocks (Supplementary Figure 7). Remarkably, overlap with both DNA methylation clocks^13,14^ and published plasma proteomic aging signatures^7,15–17^ was limited. A total of 231 of the 249 CSF clock proteins were absent from the plasma aging resources examined, indicating that the CSF clock predominantly captures molecular processes that are not represented in peripheral aging biomarkers. These findings suggest that biological brain aging is governed by a largely distinct proteomic architecture that cannot be fully inferred from blood-based aging measures. Together, all results support a model in which CSF brain-age acceleration is generated by two coordinated but opposing processes: pro-aging activation of inflammatory, vascular/BBB, ECM-remodeling and complement/coagulation programs, and anti-aging collapse of synaptic, neuronal, axonal and structural maintenance programs.

### CSF aging-clock proteins map to disease-vulnerable neurovascular and neuronal circuits across the human brain

To place the CSF aging clock in a tissue context, we projected clock-derived gene modules onto a human brain single-cell atlas comprising 2.25 million cells/nuclei, generated by our collaborators ^18^. These modules had been defined from CSF clock proteins on the basis of coefficient direction and functional reclustering, and were not re-derived from the single-cell data. Module scores were calculated using an expression-matched control-gene strategy, such that each score reflected relative enrichment of a clock-derived gene set rather than cell abundance or total expression (Fig. 6A). At the cellular level, pro-aging activation modules showed the strongest enrichment in non-neuronal and barrier-associated compartments. Vascular cells and fibroblast-like populations showed consistently high scores across inflammatory, vascular/BBB, ECM-remodeling and complement/coagulation modules. Additional pro-aging signal was observed in microglia, choroid plexus, ependymal cells and oligodendrocyte-lineage populations, indicating that the positive-coefficient component of the CSF clock mapped predominantly to neurovascular, immune, barrier and matrix-remodeling cell states (Fig. 6B). By contrast, anti-aging modules were preferentially enriched in neuronal populations. Adhesion/structural maintenance and neuronal/axonal connectivity programs were prominent in upper-layer and deep-layer excitatory neurons, hippocampal neurons and thalamic excitatory neurons, whereas synaptic/neurotransmission-related programs were enriched in inhibitory and medium spiny neuronal populations. These patterns indicate that negative-coefficient clock proteins mapped primarily to neuronal connectivity, synaptic signalling and tissue-maintenance programs rather than to the same non-neuronal compartments that carried the pro-aging signal.

**Figure 6.**
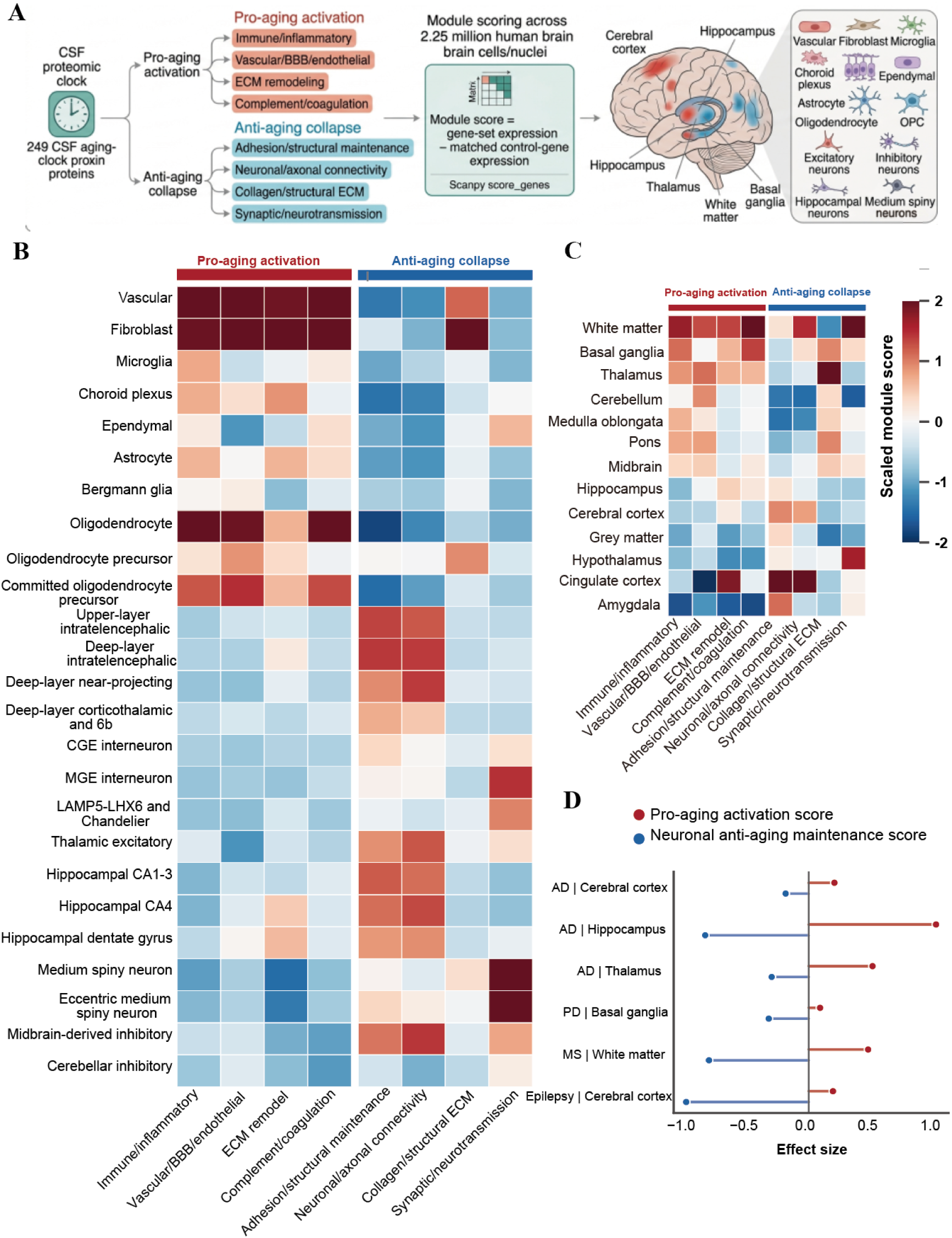
Single-cell and regional mapping of CSF aging-clock programs. **(A)** Overview of the single-cell and regional mapping framework. CSF aging-clock genes were grouped into pro-aging modules (immune/inflammatory, vascular/BBB/endothelial, extracellular matrix (ECM)-remodeling, and complement/coagulation pathways) and anti-aging modules (neuronal/axonal connectivity, synaptic/neurotransmission, adhesion/structural maintenance, and collagen/structural ECM pathways). Module scores were calculated in a human brain single-cell atlas comprising 2.25 million cells and nuclei and subsequently summarized across cell populations and anatomical brain regions. **(B)** Heatmap showing scaled module scores across major brain cell classes and neuronal subtypes. Rows represent clock-derived functional modules and columns represent cell populations. Higher values indicate stronger relative module expression. **(C)** Heatmap showing scaled module scores across anatomical brain regions. Rows represent clock-derived functional modules and columns represent brain regions. Values are z-score standardized across regions. **(D)** Effect sizes of pro-aging activation and neuronal anti-aging maintenance scores in disease-vulnerable brain regions, including Alzheimer’s disease-associated cortex, hippocampus and thalamus, Parkinson’s disease-associated basal ganglia, multiple sclerosis-associated white matter, and epilepsy-associated cerebral cortex. Positive values indicate relative enrichment, whereas negative values indicate relative depletion compared with the remaining brain regions.

Regional module scoring further showed that the two programs were anatomically organized across the human brain. Pro-aging activation scores were higher in white matter, basal ganglia, thalamus and brainstem-related regions, whereas anti-aging modules showed stronger regional structure across cortical, cingulate, hippocampal and hypothalamic regions (Fig. 6C). We next tested whether these programs were altered in disease-vulnerable regions by comparing disease samples with region-matched healthy controls. Across Alzheimer’s disease-relevant cerebral cortex, hippocampus and thalamus, Parkinson’s disease-relevant basal ganglia, multiple-sclerosis white matter and epilepsy cerebral cortex, disease samples showed a consistent shift toward higher pro-aging activation and lower neuronal anti-aging maintenance scores (Fig. 6D). The largest anti-aging reductions were observed in multiple-sclerosis white matter and epilepsy cortex, whereas Alzheimer’s disease hippocampus showed the strongest pro-aging shift. These results support a spatially resolved model in which disease-vulnerable brain compartments show activation of inflammatory, vascular and remodeling programs together with attenuation of neuronal maintenance programs.

To determine whether this composite pattern was driven uniformly by all submodules, we decomposed the disease-region analysis into individual pro-aging and anti-aging modules (Supplementary Fig. 8). In Alzheimer’s disease-relevant regions, inflammatory, vascular/BBB, ECM-remodeling and complement/coagulation modules were broadly increased, while synapse/adhesion, neuronal/axonal connectivity and synaptic/neurotransmission modules were generally reduced. Similar reductions in neuronal anti-aging modules were observed in multiple-sclerosis white matter and epilepsy cortex. However, the collagen/structural ECM module showed a distinct pattern, with increased scores in several disease-relevant regions, including Alzheimer’s disease hippocampus, Parkinson’s disease basal ganglia and multiple-sclerosis white matter. This module-level decomposition indicates that structural ECM genes capture a remodeling or stromal-response component rather than a uniformly declining neuronal maintenance program. Therefore, the primary disease-region analysis used a neuronal anti-aging maintenance score that excluded the collagen/structural ECM module, while the full module-level decomposition was retained as a sensitivity analysis.

To connect CSF clock proteins with regional brain aging, we first defined region-enriched transcriptional signatures from healthy brain pseudobulk profiles and then asked which of these signatures contained CSF aging-clock genes. A subset of clock genes showed marked regional specificity (Supplementary Fig. 9) and age-associated expression changes concordant with their CSF clock coefficients (Supplementary Fig. 10). Pro-aging genes were enriched in disease-relevant neurovascular programs, including LRG1, ICOSLG, CFI and CD14 in the basal ganglia and VWF, ESAM, CDH5, MXRA8, LTBP2 and COL6A3 in the hippocampus. In contrast, anti-aging genes showed regionally localized age-associated decreases, including NPTXR in the cingulate cortex, MET in the cerebral cortex, PCDHGC5 in the hippocampus and TFF3 in the midbrain. These results support a model in which CSF brain-age acceleration reflects spatially organized aging programs across vulnerable brain regions rather than a uniform whole-brain signal.

Single-cell localization of these region-enriched genes further linked the clock programs to specific cellular compartments (Supplementary Fig. 11). Vascular and ECM-associated genes, including HSPG2, VWF, ESAM, CDH5, MXRA8, LTBP2 and COL6A3, were localized predominantly to vascular and fibroblast-like cells in hippocampal regions. Immune and inflammatory genes, including CD14, CFI, ICOSLG and LRG1, mapped to microglial, vascular or barrier-associated compartments in the basal ganglia. In contrast, neuronal anti-aging genes showed cell-type-restricted expression in neuronal populations, including NPTXR in cingulate cortical neurons, MET in cerebral cortical neuronal populations and PCDHGC5 in hippocampal neuronal populations. Together, these analyses provide anatomical and cellular context for the CSF proteomic aging clock and indicate that CSF brain-age acceleration reflects coordinated remodeling of disease-vulnerable neurovascular, immune and neuronal networks.

### CSF proteomic aging follows a non-linear trajectory with a midlife remodeling wave and disease-specific temporal programs

Finally, we investigated when age-associated proteomic remodeling occurs during the lifespan. Sliding-window analysis revealed a highly dynamic and non-linear trajectory of CSF proteomic aging, contrary to a gradual linear aging process. Across the overall population, the proportion of age-associated proteins increased progressively from early adulthood, reached a pronounced peak around 50 years of age, and subsequently declined, forming a distinct midlife remodeling wave (Fig. 7A).

**Figure 7.**
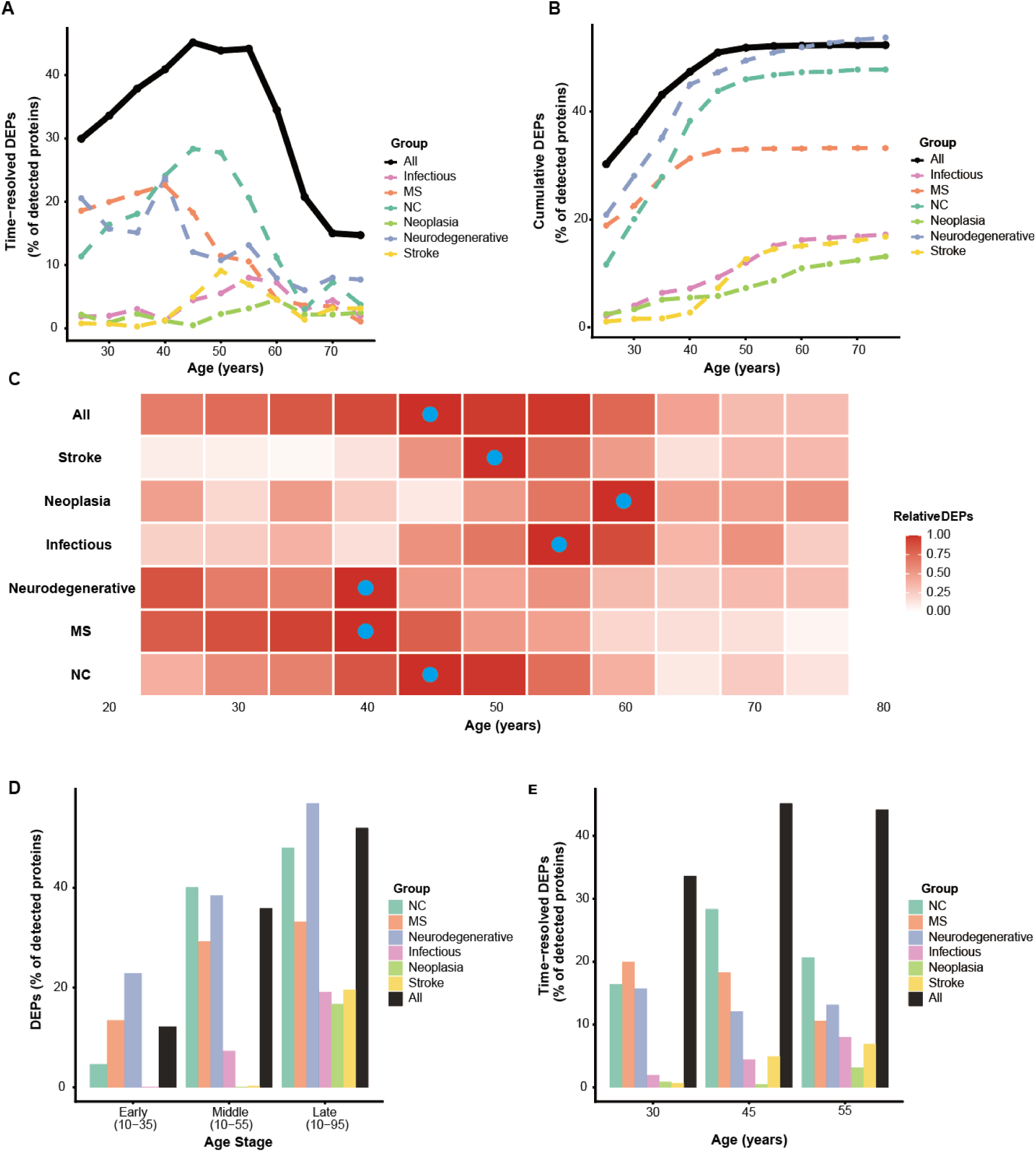
Temporal dynamics of CSF proteomic remodeling across the lifespan and neurological disease groups. **(A)** Time-resolved differentially expressed proteins (DEPs) across sliding age windows (20-year window, 5-year step), expressed as the percentage of detected proteins within each group. Curves represent the overall cohort and individual disease categories. **(B)** Cumulative accumulation of aging-associated DEPs across the lifespan. Values represent the cumulative proportion of unique DEPs identified across successive sliding windows, normalized to the number of proteins detected within each group. **(C)** Heatmap showing temporal sensitivity of age-associated proteomic remodeling across disease groups. Color intensity indicates the relative proportion of time-resolved DEPs normalized to the within-group maximum. Red circles denote the age window with the highest proportion of DEPs for each group. **(D)** Proportion of age-associated DEPs across cumulative aging stages, including Early (10–35 years), Middle (10–55 years), and Late (10–95 years) stages. Values are expressed as percentages of detected proteins within each group. **(E)** Cross-sectional summaries of age-associated proteomic remodeling at representative ages (30, 45, and 55 years). Values represent the proportion of DEPs detected within each disease category at the corresponding age. NC, neurologically normal controls; MS, multiple sclerosis.

Importantly, different neurological diseases displayed distinct temporal aging trajectories. Neurodegenerative diseases and multiple sclerosis exhibited earlier peaks of proteomic remodeling, centered around 40 years of age, whereas stroke and neoplasia showed delayed peaks occurring predominantly between 55 and 60 years (Fig. 7A). These findings suggest that accelerated brain aging does not emerge uniformly across disorders but instead follows disease-specific temporal programs that may reflect differences in underlying pathogenic mechanisms.

Cumulative analyses further demonstrated progressive accrual of aging-associated proteins across the lifespan (Fig. 7B). However, the timing and magnitude of protein accumulation differed substantially among disease groups, and temporal sensitivity maps confirmed that the most pronounced remodeling events consistently clustered within the fourth to sixth decades of life (Fig. 7C). Stage-based analyses similarly revealed increasing proportions of differentially expressed proteins from early to middle and late aging stages (Fig. 7D–E). Collectively, these observations identify midlife as a critical biological transition period during which large-scale reorganization of the CSF proteome occurs.

To understand the biological processes underlying this temporal remodeling, we examined stage-specific functional enrichment patterns. Proteins that increased during aging were consistently enriched for vascular development, angiogenesis, humoral immune responses, extracellular matrix remodeling, and lipid-associated pathways (Supplementary Fig. 12). Notably, vascular and immune programs emerged early and remained prominent across multiple aging stages, whereas lipid remodeling pathways became increasingly evident during later-life transitions.

In contrast, proteins that declined during aging were enriched for neuronal and structural maintenance processes, including synaptic organization, neuron projection development, axonogenesis, nervous system development, cell-cell adhesion, and extracellular structural integrity (Supplementary Fig.13). These pathways exhibited progressive attenuation across aging stages, suggesting cumulative erosion of neuronal resilience and connectivity over time. Together, these temporal analyses provide an independent validation of the two-direction architecture of the CSF aging clock.

## Discussion

This study establishes CSF proteomic brain-age acceleration as a quantitative and biologically interpretable measure of CNS aging. Rather than serving only as an age-prediction model, the 249-protein clock resolved a disease-relevant aging phenotype across neurological disorders. The central finding is that accelerated CSF brain aging reflects two opposing molecular programs: pro-aging activation of immune, vascular, BBB, extracellular-matrix and coagulation-related biology, and anti-aging collapse of neuronal, synaptic, axonal and structural-maintenance programs. This dual-axis structure provides a framework for interpreting why diverse neurological diseases can converge on an older-than-expected CSF proteomic state despite distinct etiologies.

The compartment in which aging was measured is central to the study. Most molecular aging clocks have been developed from blood, DNA methylation or plasma proteomics^7,13–17^. These approaches have transformed aging research, but they do not directly sample the molecular environment of the brain^19^. CSF is anatomically and physiologically closer to brain parenchyma, meninges, choroid plexus, ventricular surfaces and neurovascular interfaces^20^. The limited overlap between the CSF clock and blood-derived aging signatures therefore should not be viewed as a limitation of reproducibility. Instead, it suggests that CSF captures a CNS-enriched aging program that is partly distinct from systemic aging. This distinction is important because brain aging is unlikely to be fully inferred from peripheral biomarkers alone.

Cross-cohort validation supports the robustness of this interpretation. The clock was trained in neurologically normal controls and retained age-related structure across more than 10,000 CSF samples and three proteomic technologies, including dia-PASEF, Orbitrap-based mass spectrometry and SomaScan aptamer profiling. Prediction accuracy was strongest in neurologically normal controls and attenuated in disease-enriched or cross-platform settings, particularly after transfer to aptamer-based and TMT-based datasets. This attenuation is expected given differences in assay chemistry, protein coverage, cohort composition and calibration. For this reason, absolute predicted CSF age should be interpreted within platform-specific context, whereas calibrated age acceleration is the more meaningful biological readout. The reproducible disease-associated shifts across independent cohorts argue that CSF age acceleration is not simply a technical artifact of one platform or one diagnostic category.

The disease analyses indicate that CSF brain-age acceleration is a convergent state rather than a disease label. Increased AgeAccel was observed across inflammatory, infectious, neoplastic, vascular and neurodegenerative conditions, and rose progressively across the Alzheimer’s disease continuum. However, the same age-accelerated state was not produced by identical protein changes in every disease. SHAP and disease-category analyses suggest that different disorders perturb partially distinct combinations of clock proteins, with immune and barrier-associated proteins more prominent in inflammatory conditions, vascular and injury-related proteins in stroke-associated contexts, and neuronal or structural-maintenance losses in neurodegenerative settings. This distinction matters because it prevents overinterpretation of AgeAccel as a diagnostic classifier. CSF AgeAccel is better understood as a state variable that integrates the extent to which a disease has shifted the CSF proteome away from a normal aging trajectory.

The pro-aging axis was biologically coherent and clinically relevant. Representative proteins such as CHI3L1, CD14, LRG1, VWF and LTBP2 connect accelerated CSF aging to glial and innate immune activation, endothelial dysfunction, vascular remodeling, BBB permeability and extracellular-matrix turnover. These proteins are not redundant markers of generic inflammation. VWF provides a vascular and endothelial link, CD14 and CHI3L1 capture innate immune and glial activation, LRG1 connects vascular-inflammatory remodeling, and LTBP2 reflects matrix and barrier-associated structural change^21–25^. Functional grouping further showed enrichment of immune/inflammatory, vascular/BBB/endothelial, ECM-remodeling and complement/coagulation modules. Together, these findings suggest that the pro-aging component of the clock reflects coordinated activation of barrier-vascular-matrix biology, a process increasingly recognized as central to brain aging and neurological vulnerability.

The anti-aging axis is equally important. Proteins including NPTX2, COL1A2, NID1, CDH8, PENK and CNTN3 link lower-than-expected CSF aging to synaptic maintenance, neuronal connectivity, cell adhesion, neuropeptide signaling and structural extracellular-matrix integrity^26–31^. Their decline with higher AgeAccel indicates that accelerated CSF aging is not merely the accumulation of inflammatory or vascular injury signals. It also reflects loss of proteins associated with preserved neuronal architecture, synaptic resilience and tissue structural support. This anti-aging collapse provides a useful counterweight to inflammation-centered models of brain aging. It separates two processes that may occur together but are not identical: activation of damage-associated neurovascular programs and withdrawal of neuronal, synaptic and structural-maintenance programs.

The BBB findings provide a physiological bridge between the pro-aging axis and clinical CSF measurements. AgeAccel was higher in individuals with elevated QAlb and increased stepwise across QAlb tertiles. Additional CSF barrier and intrathecal-immunity measures, including QIgG, CSF albumin, CSF IgG and oligoclonal bands, were positively associated with AgeAccel. These results place BBB dysfunction close to the biological core of the clock. However, the relationship remains associative. Barrier dysfunction may contribute to the aging signature by altering protein entry, immune exposure and vascular-matrix remodeling, but it may also reflect disease severity or inflammatory tissue injury.

The longitudinal ADNI analyses extend the clock from a cross-sectional aging marker to a prognostic phenotype. Higher baseline AgeAccel predicted faster deterioration across clinical, fluid biomarker, MRI and PET outcomes and was associated with increased risk of diagnostic progression from MCI to dementia and from normal cognition to cognitive impairment. Importantly, a reduced Top30 Protein Panel retained much of this prognostic directionality. This finding is translationally important because a 249-protein mass-spectrometry clock is powerful for discovery but may be difficult to implement as a routine clinical assay. A smaller panel that samples both axes of the clock could enable more feasible targeted proteomic or immunoassay validation. Such a panel should not be selected only from the largest positive coefficients. It should preserve both vascular-inflammatory activation and neuronal-maintenance decline, allowing patients to be stratified into pro-aging-dominant, anti-aging-collapse-dominant or mixed profiles.

Single-cell and regional mapping provided anatomical context for the two-axis model. Module scores across 2.25 million human brain cells/nuclei showed that pro-aging programs were enriched in non-neuronal and barrier-associated compartments, including vascular, fibroblast, microglial, choroid plexus and oligodendrocyte-lineage populations. In contrast, anti-aging modules were enriched in neuronal populations and regionally specialized neuronal subtypes, including cortical, hippocampal, thalamic, inhibitory and medium spiny neuronal populations. Brain-region analyses further linked these programs to disease-vulnerable regions, including Alzheimer’s disease-associated cortex, hippocampus and thalamus, Parkinson’s disease-associated basal ganglia, multiple sclerosis-associated white matter and epilepsy-associated cerebral cortex. These analyses do not prove that the corresponding cells are the direct source of CSF proteins, because transcript abundance, protein abundance, secretion and CSF detectability are not equivalent. Nevertheless, they provide a plausible tissue framework in which CSF AgeAccel reflects coordinated remodeling of neurovascular, glial and neuronal compartments.

The genetic analyses add an additional layer of disease specificity. Parkinson’s disease GWAS signals showed nominal enrichment in the full clock, positive-coefficient genes, immune/inflammatory modules, vascular/BBB/endothelial modules and neuronal/axonal connectivity modules. Direct overlap included BST1, STX1B, GPNMB, EFNA3 and PRSS8, all belonging to the positive-coefficient pro-aging axis. By contrast, Alzheimer’s disease GWAS overlap was limited and did not show comparable enrichment across clock-derived gene sets. They suggest that CSF aging-clock biology converges more strongly with Parkinson’s disease genetic susceptibility than with Alzheimer’s disease genetic association signals in the current analysis, particularly through immune-vascular and neuronal-connectivity pathways. They do not imply that CSF AgeAccel is specific to Parkinson’s disease, nor do they exclude non-genetic or downstream disease processes in Alzheimer’s disease.

The temporal analyses suggest that CSF proteomic aging is not a linear accumulation of changes across the lifespan. Sliding-window and stage-based analyses identified a midlife remodeling wave, with prominent proteomic changes around the fourth to sixth decades and disease-specific timing of peak remodeling. Upregulated proteins during aging were enriched for vascular development, angiogenesis, immune response, ECM remodeling and lipid-associated pathways, whereas downregulated proteins were enriched for synaptic organization, neuron projection development, axonogenesis, cell adhesion and structural integrity. This temporal architecture independently supports the two-axis model: vascular-inflammatory and matrix programs rise as neuronal and synaptic maintenance programs decline. Because these analyses are cross-sectional, they identify an age window of heightened remodeling but do not establish that midlife remodeling causes later neurological disease. Longitudinal midlife cohorts will be needed to test this possibility.

Several limitations define the boundaries of the present conclusions. First, many analyses were cross-sectional, so temporal order and causality cannot be inferred. Second, diagnostic groups were heterogeneous and may differ in disease duration, severity, treatment exposure, comorbidity and CSF sampling indication. Third, cross-platform application required calibration and was affected by differences in protein coverage and assay chemistry. Fourth, CSF integrates signals from brain parenchyma, meninges, choroid plexus, vasculature, immune compartments and blood-derived proteins, making precise cellular origin difficult to assign. Fifth, single-cell mapping provides anatomical plausibility but not direct protein-source validation. Finally, BBB-associated proteins may partly encode permeability or blood-derived protein entry, so sensitivity analyses excluding or adjusting for barrier-related proteins will be important.

Future work should move in three directions. First, longitudinal CSF cohorts should test whether AgeAccel precedes cognitive decline, Parkinson’s disease progression, multiple sclerosis disability, post-stroke recovery or treatment response. Second, targeted proteomics or immunoassays should validate whether a compact pro-aging and anti-aging panel can reproduce the biological and prognostic information of the full clock. Third, mechanistic studies should determine whether modifying BBB integrity, vascular remodeling, inflammation or synaptic maintenance alters the CSF aging score. These steps would move the CSF clock from a discovery framework toward a testable biological and translational tool.

In summary, CSF proteomic brain-age acceleration defines a two-axis phenotype of human CNS aging. It increases across neurological diseases, predicts longitudinal deterioration, and reflects the combined effect of pro-aging neurovascular and inflammatory activation with anti-aging neuronal and synaptic maintenance loss. By measuring both sides of this process, the CSF aging clock provides a framework for quantifying biological brain aging beyond systemic aging markers and for developing clinically tractable CSF panels that capture disease-relevant CNS vulnerability.

## Methods

### Human cohorts

#### TimsTOF Cohort

The primary discovery cohort comprised 6,189 cerebrospinal fluid (CSF) samples collected from patients undergoing diagnostic lumbar puncture at the Department of Neurology, Technical University of Munich (TUM), Germany, as previously described^12^. Clinical and proteomic measurements were conducted in a blinded fashion. The cohort comprised 1,535 individuals with multiple sclerosis, 1,518 with other CNS diseases, 856 with other neurological autoimmune diseases, 841 neurologically normal controls, 497 with neurodegenerative diseases, 437 with infectious diseases, 306 with neoplasia, and 199 with ischemic stroke. Diagnoses were extracted from hospital records and made according to standard clinical guidelines and diagnostic criteria at the time of sample collection. The control group consisted of individuals who received diagnostic workup after presenting with headache with no evidence of structural neurological disease or a diagnosis of idiopathic intracranial hypertension. MS diagnoses were made according to the McDonald criteria valid at the time of diagnosis.

All CSF samples were measured on the timsTOF Pro 2 mass spectrometer (Bruker Daltonics) coupled to an Evosep One chromatography system. Samples were separated using a 21-minute gradient with a flow rate of 1 μl/min during peptide elution, corresponding to a throughput of 60 samples per day. The mass spectrometer was operated in dia-PASEF mode with one MS1 scan followed by ten dia-PASEF scans covering 20 MS2 windows. An optimized dia-PASEF method based on the precursor density distribution in the m/z – ion mobility plane was applied, with constrains of m/z 400–1200 and ion mobility 0.8–1.3 Vs cm⁻² 1/K₀. All samples were processed and analyzed with uniform quality control standards. Written informed consent was obtained from all individuals, and the local ethics committee approved the study (S324/2019, TUM).

#### Astral-1 Cohort

A stratified discovery subset of 1,965 CSF samples was selected from the main discovery cohort and measured on the Orbitrap Astral mass spectrometer (Thermo Fisher Scientific) to enhance biomarker discovery for multiple sclerosis diagnosis. The cohort comprised 978 individuals with MS, 427 with other neurological autoimmune diseases, 221 neurologically normal controls, 219 with infectious diseases, 85 with other CNS diseases, 13 with neoplasia, 11 with neurodegenerative diseases, and 11 with ischemic stroke. Sample stratification was performed by clinical specialists to reduce covariate differences and retain only samples with the clearest and most relevant diagnoses. Inclusion criteria included minimal blood contamination (0 or (+) erythrocytes), with stricter requirements for MS samples (no evidence of blood contamination). All samples were measured on the Orbitrap Astral mass spectrometer coupled to an EvoSep One chromatography system with an 11.5-minute gradient, corresponding to 100 samples per day throughput.

#### Astral-2 Cohort

An independent replication cohort of 615 new CSF samples was collected from the same institution using identical sample collection and processing protocols. The cohort comprised 219 individuals with MS, 204 with other autoimmune diseases, 155 neurologically normal controls, and 37 with infectious diseases. Inclusion criteria required available information on QAlb, blood contamination status, and OCB status. All samples in the Astral-2 replication cohort were measured on the Orbitrap Astral platform using identical parameters to the Astral-1 discovery subset, enabling independent external validation of the proteomic aging clock.

#### PPMI cohort

An independent external validation cohort was obtained from the Parkinson’s Progression Markers Initiative (PPMI), an international observational study designed to identify biomarkers of Parkinson’s disease progression. The analyzed cohort comprised 1,153 CSF samples, including 969 individuals with Parkinson’s disease and 184 healthy controls. CSF proteomics was measured using the SomaLogic SomaScan 4k platform, which quantifies protein analytes using modified single-stranded DNA aptamers. Protein abundance values were provided as batch-corrected, log2-transformed relative fluorescence units from PPMI Project 151. SOMAmer identifiers were mapped to HGNC gene symbols using SomaLogic annotation files; for analytes targeting multiple proteins, the SOMAmer identifier was retained to preserve feature uniqueness.

Clinical and demographic data were obtained from the PPMI curated data release and merged with proteomic data using participant ID and visit code. To apply the timsTOF-trained CSF aging clock to SomaScan data, overlapping clock proteins were identified by gene-symbol matching. Because of platform differences between dia-PASEF mass spectrometry and aptamer-based proteomics, predicted ages were calibrated within healthy controls by z-score remapping: predicted ages were standardized to the mean and standard deviation of healthy-control predicted ages and then rescaled to the mean and standard deviation of healthy-control chronological ages. AgeAccel was defined as the residual from a linear regression of calibrated predicted age on chronological age.

#### ADNI Cohort

An independent external validation cohort was obtained from the Alzheimer’s Disease Neuroimaging Initiative (ADNI), a multicentre longitudinal study designed to develop clinical, imaging, genetic and biochemical biomarkers for Alzheimer’s disease. The analyzed cohort comprised 1,105 baseline CSF samples, including 379 cognitively normal controls, 562 individuals with mild cognitive impairment and 164 individuals with Alzheimer’s disease dementia. CSF proteomics was measured by Emory University using tandem mass tag mass spectrometry (TMT-MS), as described in the ADNI Emory CSF TMT-MS dataset. CSF samples were digested, labelled with TMTpro 18-plex reagents, fractionated by high-pH reversed-phase chromatography and analyzed by LC–MS/MS with FAIMS Pro ion mobility separation. Protein searches were performed using FragPipe against the canonical human UniProt proteome. A total of 3,447 proteins were quantified across baseline CSF samples.

Protein identifiers were provided as HGNC gene symbol concatenated with UniProt accession. Gene symbols were extracted from the prefix before the first underscore, and duplicate gene entries were resolved before model application to retain a single feature per gene. Values coded as zero or −4 in the released matrix were treated as missing where appropriate. Clinical and demographic variables, including age, sex, education and APOE ε4 status, were obtained from the ADNI repository and merged with proteomic data using participant identifier and visit code. Diagnostic groups were defined using ADNI clinical diagnosis fields as cognitively normal, mild cognitive impairment or Alzheimer’s disease dementia.

To apply the timsTOF-trained clock to ADNI TMT-MS data, model features not quantified in ADNI were imputed using the corresponding training-set mean values stored in the model scaling parameters. Feature-wise distribution alignment was then applied to reduce systematic platform differences: each ADNI protein feature was standardized using its own mean and standard deviation and rescaled to the corresponding training-set mean and standard deviation. The locked elastic-net model was then applied without retraining. Predicted ages were calibrated within cognitively normal controls using rank-based quantile mapping, aligning the empirical distribution of predicted age to the chronological-age distribution in controls while preserving rank ordering across participants. ADNI AgeAccel was defined as the residual from a linear regression of calibrated predicted age on chronological age across all ADNI samples.

### Model Training and Selection

To construct the CSF proteomic aging clock, regularized regression models were trained to predict chronological age from CSF protein abundance profiles in neurologically normal controls from the timsTOF discovery cohort (n = 841). Ridge regression, LASSO and elastic-net models were compared across α values from 0 to 1 in increments of 0.1 using the glmnet package in R. For each α, λ was selected by tenfold cross-validation, and λ.1se was used to favour model sparsity and generalizability. The final model was selected based on cross-validated mean squared error. The optimal model was an elastic net with α = 0.1, yielding a 249-protein CSF aging clock that predicted chronological age in discovery controls with high accuracy (r = 0.937, MAE = 3.82 years).

### Definition of CSF age acceleration

Predicted CSF age was obtained by applying the locked aging-clock model to standardized protein abundance profiles. Raw age gap was calculated as predicted CSF age minus chronological age. The primary age-acceleration metric was defined as the residual from a linear regression of predicted or calibrated predicted CSF age on chronological age. For cross-platform external cohorts, calibrated predicted age was used before deriving AgeAccel. Sensitivity age-acceleration metrics were additionally derived from residual models adjusting for sex, sample-preparation batch, QAlb or CSF leukocyte count where these covariates were available.

### Disease-group and statistical analyses

Disease-associated CSF AgeAccel was assessed by comparing each diagnostic group with neurologically normal controls within the corresponding cohort. Two-sided Wilcoxon rank-sum tests were used for group comparisons. In discovery and Astral validation analyses, Benjamini–Hochberg correction was applied across diagnostic comparisons within each analysis. Sex-stratified analyses were performed by repeating disease-versus-control comparisons separately in male and female participants. In PPMI, Parkinson’s disease and healthy controls were compared after healthy-control-based calibration. In ADNI, cognitively normal controls, mild cognitive impairment and Alzheimer’s disease dementia were compared using two-sided Wilcoxon rank-sum tests.

### Clock-protein directionality, contribution and disease heatmaps

Clock proteins were classified by the sign of their elastic-net coefficients. Proteins with positive coefficients were defined as pro-aging proteins, indicating that higher standardized abundance contributed to older predicted CSF age. Proteins with negative coefficients were defined as anti-aging proteins, indicating that higher standardized abundance contributed to younger predicted CSF age. Spearman correlations between individual clock proteins and AgeAccel were calculated to evaluate concordance between coefficient-defined directionality and observed age-acceleration-associated abundance patterns.

Protein-level contribution scores were calculated by multiplying each standardized protein abundance value by its corresponding model coefficient. Group-level contribution profiles were obtained by averaging coefficient-weighted contributions within each diagnostic group. For disease-specific heatmaps, selected clock proteins were compared between each neurological disease group and neurologically normal controls. Log2 fold changes relative to controls were visualized, and marked proteins were those overlapping pathway-defined core proteins from the pro-aging and anti-aging module analyses.

### Functional reclustering of pro-aging and anti-aging proteins

To interpret the biological architecture of the clock, positive- and negative-coefficient proteins were analyzed separately. Protein lists were submitted for functional enrichment and protein-network annotation using STRING and Gene Ontology biological process terms. Enriched terms and their contributing proteins were manually consolidated into biologically related mechanism classes, guided by term similarity, protein membership and known CNS biology. Pro-aging proteins were grouped into immune/inflammatory, vascular/BBB/endothelial, ECM-remodeling and complement/coagulation modules. Anti-aging proteins were grouped into synaptic junction/adhesion, neuronal/axonal connectivity, collagen/structural ECM and neuropeptide-transmission modules. Module assignments were used for visualization, gene-set scoring and GWAS enrichment analyses. Because some proteins participate in more than one biological process, functional interpretation was based on module-level patterns rather than assuming strictly mutually exclusive protein functions.

### BBB permeability and intrathecal immune marker

Blood–brain barrier permeability was assessed using the CSF/serum albumin quotient (QAlb). Elevated QAlb was defined using age-specific thresholds: QAlb > 6.5 for participants younger than 40 years, QAlb > 8.0 for participants aged 40–59 years and QAlb > 10.0 for participants aged 60 years or older. QAlb tertiles were generated within the analyzed cohort. AgeAccel differences between QAlb groups were tested using two-sided Wilcoxon rank-sum tests.

Associations between AgeAccel and CSF barrier or intrathecal immune markers were assessed using age- and sex-adjusted partial Spearman correlations. Markers included QIgG, QAlb, CSF albumin, CSF IgG and oligoclonal bands. Correlation coefficients and 95% confidence intervals were reported, and multiple testing was controlled using Benjamini–Hochberg false-discovery-rate correction.

### Reduced 30-protein panel

The Top30 Protein Panel was generated by selecting the 30 proteins with the largest absolute coefficients from the full 249-protein CSF aging clock. A reduced age-prediction model was refitted in neurologically normal discovery controls using these 30 proteins and then applied to ADNI using the same preprocessing, feature-alignment and calibration workflow used for the full model. Predicted ages were calibrated using cognitively normal ADNI participants, and AgeAccel was calculated as the residual of calibrated predicted age after regression on chronological age. AgeAccel values were z-standardized before longitudinal and survival analyses.

### ADNI longitudinal outcomes and diagnostic progression

Baseline ADNI AgeAccel was tested for association with longitudinal clinical, fluid biomarker, MRI and PET outcomes. Continuous longitudinal outcomes were transformed into worsening-oriented z scores so that higher values consistently represented greater deterioration. Linear mixed-effects models were used to test whether baseline AgeAccel modified the rate of longitudinal change. Models included baseline AgeAccel, follow-up time, the AgeAccel × time interaction, chronological age, sex and education as fixed effects, with a participant-specific random intercept. The interaction term was interpreted as the annual worsening effect associated with higher baseline AgeAccel.

Time-to-event analyses were performed using Cox proportional hazards models for progression from mild cognitive impairment to all-cause dementia, mild cognitive impairment to Alzheimer’s disease dementia, normal cognition to cognitive impairment and normal cognition to dementia. Follow-up time was calculated from baseline CSF proteomic assessment to the first visit at which the target diagnostic transition was recorded. Participants without an event were censored at their last available diagnostic follow-up. Models included baseline AgeAccel, chronological age, sex and education. Hazard ratios were reported per 1-s.d. higher baseline AgeAccel. Kaplan–Meier curves were generated after stratifying participants into tertiles of baseline AgeAccel, and group differences were assessed using log-rank tests.

### Regional pseudobulk analysis of CSF aging-clock genes

To identify brain-region-enriched CSF aging-clock genes, pseudobulk profiles were generated by aggregating raw counts within each brain region and sample. Differential expression analysis was performed using edgeR (v4.0.1) ^32^. For each brain region, predefined CSF aging-clock genes were compared between the target region and all remaining regions. TMM normalization was applied within edgeR, and genes with a false-discovery rate (FDR) < 0.05 and log2 fold change > 1 were considered region-enriched. Regional specificity was visualized using row-wise z-scores calculated from mean TMM-normalized counts per million (CPM) values across brain regions.

To link regional specificity with aging-related transcriptional changes, Pearson correlation analysis was performed between donor chronological age and TMM-normalized gene expression within each brain region. Samples with ambiguous age annotations or donor age >95 years were excluded. P values were adjusted using the Benjamini-Hochberg procedure. Candidate region-specific clock genes were prioritized when their age-correlation direction was concordant with the corresponding CSF clock coefficient direction and the absolute Pearson correlation coefficient was >0.3.

### Single-cell module scoring and disease-region comparisons

Previously defined CSF aging-clock gene modules were projected onto the human brain single-cell atlas using sc.tl.score_genes in Scanpy (v1.10) ^33^. Scoring was performed after library-size normalization and log1p transformation. Module scores were calculated using expression-matched control genes and therefore represented relative enrichment of each clock-derived gene set rather than cell abundance or total expression. Scores were aggregated at the donor, sample, brain-region and cell-type levels for downstream analyses.

For disease-region analyses, module scores in disease samples were compared with region-matched healthy controls. Effect sizes were calculated as Cohen’s d, and statistical significance was assessed using two-sided Welch’s t-tests followed by Benjamini-Hochberg correction. The pro-aging activation score was calculated from immune/inflammatory, vascular/BBB/endothelial, ECM-remodeling and complement/coagulation modules. The neuronal anti-aging maintenance score was calculated from anti-aging modules related to adhesion/structural maintenance, neuronal/axonal connectivity and synaptic/neurotransmission programs. The collagen/structural ECM module was analysed separately in module-level decomposition because it showed disease-associated behaviour distinct from the neuronal anti-aging modules.

### MAGMA gene-set enrichment of CSF aging-clock proteins

MAGMA gene-level and competitive gene-set enrichment analyses were performed to assess whether CSF aging-clock proteins were enriched for genetic association signals in Parkinson’s disease and Alzheimer’s disease GWAS. Parkinson’s disease summary statistics were obtained from the Kim et al^34^. trans-ancestry PD GWAS meta-analysis excluding 23andMe data. Alzheimer’s disease summary statistics were obtained from the Bellenguez et al. 2022 GWAS^35^. SNP-to-gene annotation was performed using MAGMA v1.10 with the 1000 Genomes European reference panel and NCBI37.3 gene locations. For the AD GWAS, SNP P values were supplied using rsIDs and per-SNP total sample sizes calculated as cases plus controls. For the PD GWAS, variants were mapped to rsIDs using chromosome and base-pair positions in the 1000 Genomes European PLINK BIM file; 8,395,011 of 8,938,152 positions were mapped and retained. PD gene-level analysis used the reported total sample size of 1,028,993 individuals. Clock proteins were mapped from gene symbols to Entrez IDs using the NCBI37.3 gene-location file. Gene sets included the full 249-protein clock, positive- and negative-coefficient clock genes, the Top30 Protein Panel, and functionally reclustered pro-aging and anti-aging modules. Competitive gene-set enrichment was tested using one-sided positive MAGMA gene-set analysis, conditioning on default internal covariates including gene size, gene density and minor-allele-count properties. P values shown in the figure are nominal MAGMA competitive gene-set enrichment P values.

### Temporal proteomic remodeling analyses

Age-associated proteomic remodeling was analyzed in neurological controls, MS, neurodegenerative disease, infectious disease, neoplasia, stroke, and the full cohort. Lifespan-wide aging-related differentially expressed proteins were identified by Spearman correlation between protein abundance and chronological age. Proteins with Benjamini-Hochberg adjusted p < 0.05 and absolute Spearman rho > 0.1 were defined as lifespan-wide aging-related proteins and classified as increasing or decreasing with age according to the sign of rho.

Stage-specific aging analyses were performed within cumulative age ranges of 10-35 years, 10-55 years, and 10-95 years. Within each stage and diagnostic group, Spearman correlations between protein abundance and age were calculated, and proteins with adjusted p < 0.05 and absolute rho >

0.1 were considered stage-specific aging-related proteins. To capture higher-resolution temporal dynamics, a 20-year sliding window with a 5-year step was applied across the available age range. Windows with fewer than 15 samples were excluded. Within each window, lifespan-wide aging-related proteins were retested for correlation with age, and proteins with nominal p < 0.05 were defined as time-resolved aging-related proteins for that window. Counts of time-resolved proteins were normalized to the number of detected proteins in each group, and cumulative counts were calculated from unique proteins detected across successive windows. Temporal functional enrichment was performed by intersecting stage-specific aging-related proteins with lifespan-wide aging-related proteins and testing Gene Ontology biological process enrichment for each group, stage, and direction. Gene lists with fewer than five proteins were not analyzed. Enrichment p values were adjusted using the Benjamini-Hochberg method.

## Ethics Approval

The timsTOF and Astral cohort were conducted in accordance with the Declaration of Helsinki. Written informed consent was obtained from all participants, and study procedures were approved by the relevant institutional ethics committees, including the local ethics committee at TUM for the primary discovery cohort. CSF collection was performed as part of routine clinical care or approved research protocols.

PPMI data use was approved by the PPMI Steering Committee and the institutional review boards of all participating sites. Data access was granted under an approved data-use agreement. ADNI data use was approved under ADNI Data Sharing and Publications Committee procedures. ADNI was approved by the institutional review boards of all participating sites, and written informed consent was obtained from all participants or their authorized representatives.

## Data Availability

The timsTOF and Astral cohort proteomics data analyzed in this study were generated as part of a large-scale CSF proteomics initiative and are available through the PRIDE proteomics data repository^12^. Raw mass spectrometry files and processed protein abundance matrices have been deposited to the ProteomeXchange Consortium via the PRIDE partner repository. For specific data access inquiries, researchers should refer to the data availability statement of the original publication.

PPMI proteomics data (Project 151, SomaLogic SomaScan) are available through the PPMI data access procedure at https://www.ppmi-info.org/access-data-specimens/download-data. Researchers are required to complete a data use agreement and submit a research proposal reviewed by the PPMI data access committee. For further information, contact the PPMI Biorepository at

ADNI proteomics data (Emory University CSF TMT-MS) are available through the ADNI data sharing infrastructure at https://adni.loni.usc.edu via the LONI Image and Data Archive (IDA). Researchers are required to complete a data use agreement and submit an access application reviewed by the ADNI Data Sharing and Publications Committee.

Summary-level results, aggregated group statistics, and the aging clock model coefficients supporting the findings of this study are available from the corresponding author upon reasonable request, subject to institutional data sharing agreements and ethical approval.

The brain single-cell transcriptomic data analyzed in this study were obtained from the publicly available Brain Cell Atlas resource. The Brain Cell Atlas web portal and associated datasets are available at https://www.braincellatlas.org and https://www.braincellatlas.org/dataSet. Original data accessions and source repositories are provided in the corresponding Brain Cell Atlas publication.

## Code Availability

Code used for data preprocessing, CSF aging-clock application, cross-platform calibration, statistical analyses and figure generation will be made available at [GitHub repository URL] upon publication. Analyses were performed using R and Python. R packages included edgeR, glmnet, tidyverse, ggplot2, survival, lme4, ggpubr and writexl; Python analyses included Scanpy for single-cell module scoring. MAGMA v1.10 was used for GWAS gene-level and gene-set enrichment analyses. Package versions and session information are documented in the code repository.

## Acknowledgements

We thank all participants and their families who contributed samples, clinical data and biomarker data to this study. We are grateful to the clinical and technical staff at the Department of Neurology, Technical University of Munich (TUM), Germany, for sample collection, processing and clinical phenotyping.

Data used in the preparation of this article were obtained from the Parkinson’s Progression Markers Initiative (PPMI) database (https://www.ppmi-info.org/access-data-specimens/download-data). For up-to-date information on the study, visit https://www.ppmi-info.org. PPMI is a public-private partnership funded by the Michael J. Fox Foundation for Parkinson’s Research and funding partners. The PPMI CSF proteomics dataset used in this study was generated using the SomaLogic SomaScan platform.

Data collection and sharing for this project were supported by the Alzheimer’s Disease Neuroimaging Initiative (ADNI). ADNI data are disseminated by the Laboratory for Neuro Imaging at the University of Southern California. Data collection and sharing for ADNI have been funded by the National Institute on Aging, the National Institute of Biomedical Imaging and Bioengineering, the Canadian Institutes of Health Research, private-sector contributions facilitated by the Foundation for the National Institutes of Health, and additional ADNI funding partners. The ADNI CSF TMT-MS proteomics dataset was generated by the Emory University proteomics team and obtained from the ADNI repository.

## Author contributions

Z.L. supervised the work and contributed to study design, data interpretation and manuscript revision. S.X. and Z.L. designed the study. S.X. had full access to the data and carried out the statistical analyses, and takes responsibility for the integrity of the data and the accuracy of the analyses. Y.Y. and Y.G. provided clinical data, contributed to the integration and interpretation of clinical and molecular findings, and secured funding support. Z.M. and K.F. performed the single cell analyses. S.L., T.W., Y.L., M.Z., H.L. and T.X. contributed demographic, clinical, biomarker and neuroimaging data, assisted with data curation and interpretation of the results, and critically reviewed the manuscript. All authors reviewed and approved the final manuscript.

## Supplementary Figure

**Supplementary Figure 1.**
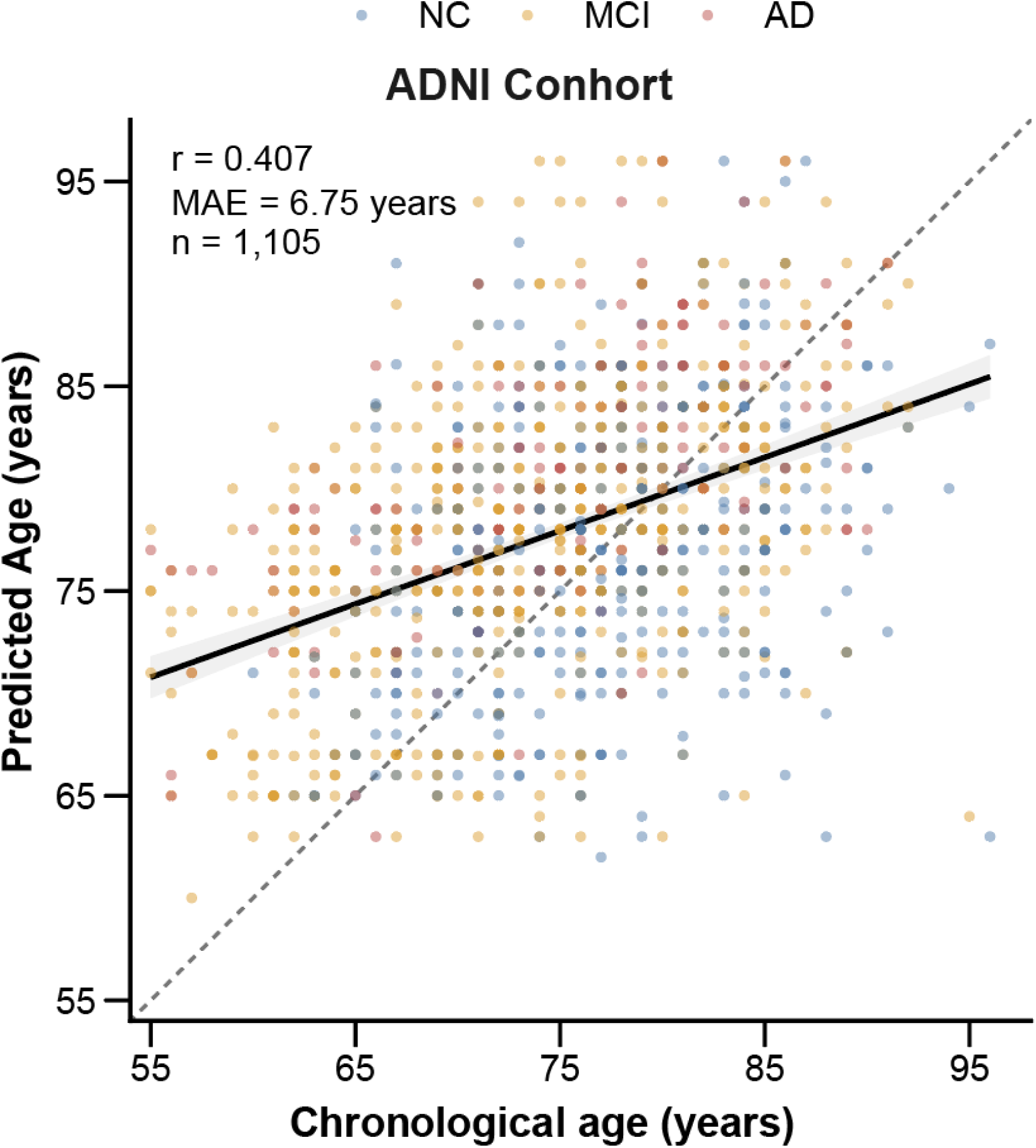
External validation of the CSF proteomic aging clock in ADNI. Scatter plot of predicted CSF proteomic age versus chronological age in the ADNI cohort (*n* = 1,105). Participants are colored according to diagnostic group (NC, MCI, and AD). The dashed line denotes the line of identity. The solid line represents the fitted linear regression, with shading indicating the 95% confidence interval. Pearson correlation coefficient (*r*) and mean absolute error (MAE) are shown.

**Supplementary Figure 2.**
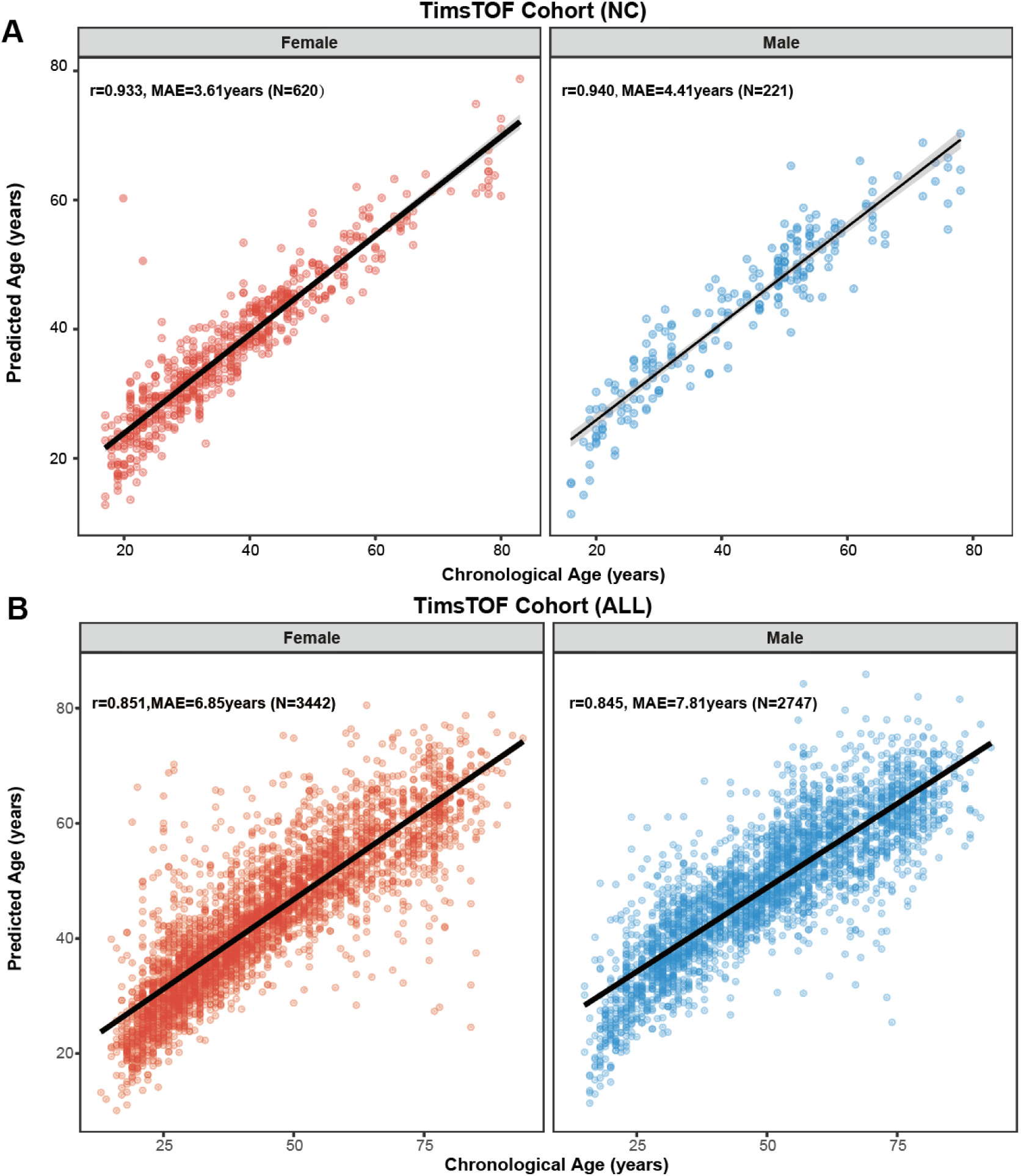
Sex-stratified performance of the CSF proteomic aging clock in the timsTOF cohort. **(A)** Age-prediction performance of the CSF proteomic aging clock in neurologically normal controls from the timsTOF cohort, stratified by sex. Scatter plots show predicted CSF proteomic age versus chronological age in females (*n* = 620) and males (*n* = 221). Pearson correlation coefficients (*r*) and mean absolute errors (MAE) are indicated in each panel (females: *r* = 0.933, MAE = 3.61 years; males: *r* = 0.940, MAE = 4.41 years). **(B)** Sex-stratified age-prediction performance in the full timsTOF cohort, including neurologically normal controls and neurological disease cases. Scatter plots show predicted CSF proteomic age versus chronological age in females (*n* = 3,442) and males (*n* = 2,747). Pearson correlation coefficients (*r*) and mean absolute errors (MAE) are shown in each panel (females: *r* = 0.851, MAE = 6.85 years; males: *r* = 0.845, MAE = 7.81 years). Each point represents one CSF sample. Solid black lines indicate linear regression fits, and gray shading denotes the 95% confidence interval. MAE, mean absolute error.

**Supplementary Figure 3.**
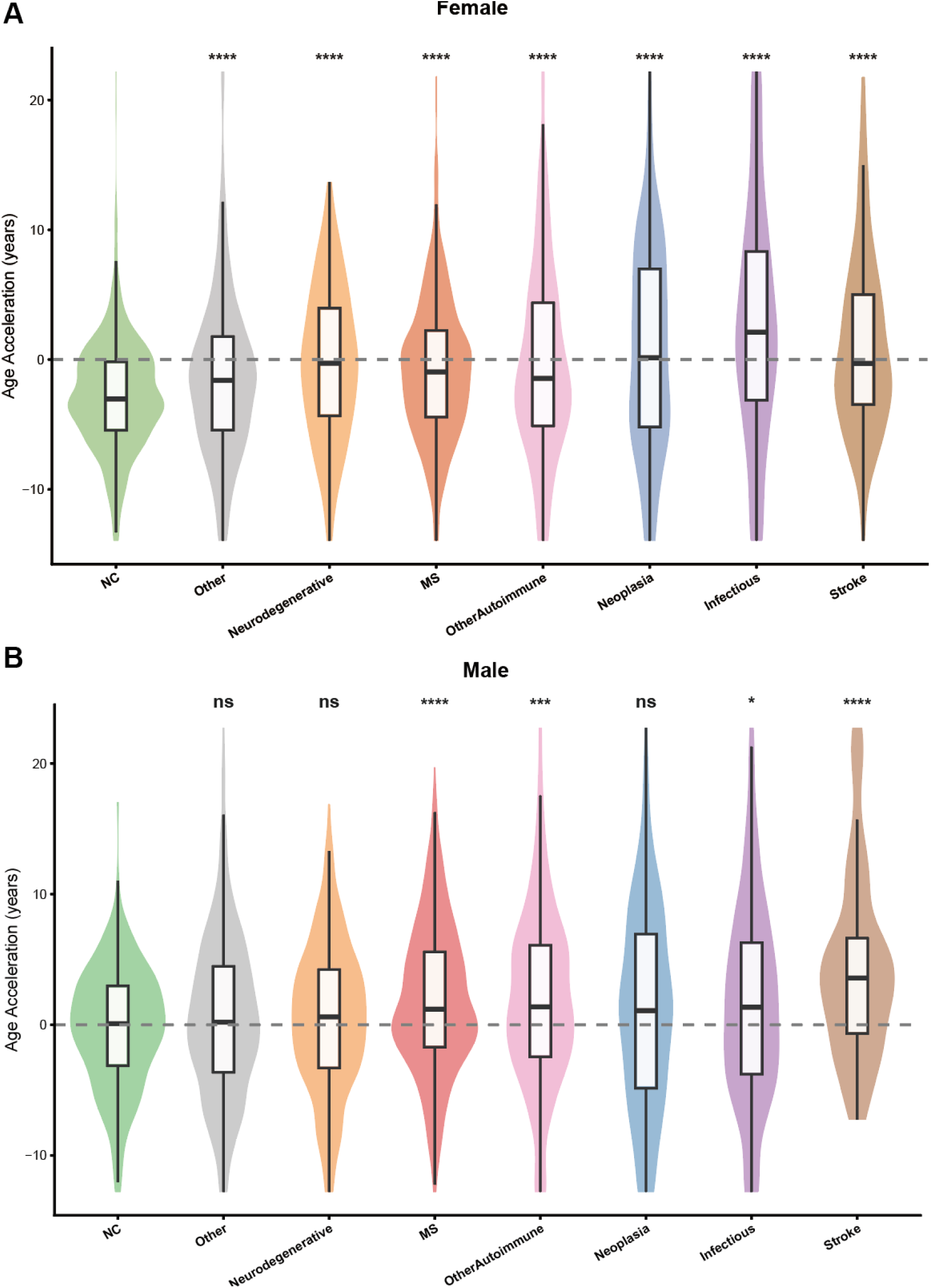
Sex-stratified CSF proteomic age acceleration across neurological disease groups. **(A, B)** Violin plots with embedded boxplots showing CSF proteomic age acceleration (AgeAccel) across diagnostic groups in female (A) and male (B) participants. AgeAccel was defined as predicted CSF proteomic age minus chronological age. Positive values indicate older-than-expected CSF proteomic age relative to chronological age, whereas negative values indicate younger-than-expected CSF proteomic age. The dashed horizontal line denotes zero age acceleration. Diagnostic groups include neurologically normal controls (NC), other neurological diseases, neurodegenerative diseases, multiple sclerosis (MS), other autoimmune neurological diseases, neoplasia, infectious diseases, and stroke. Violin plots represent the distribution of AgeAccel values within each diagnostic category. Embedded boxplots show the median (center line), interquartile range (box), and 1.5× interquartile range (whiskers). Statistical significance was assessed relative to sex-matched neurologically normal controls using two-sided Wilcoxon rank-sum tests. Significance levels are indicated as follows: ns, not significant; *P < 0.05; **P < 0.01; ***P < 0.001; ****P < 0.0001.

**Supplementary Figure 4.**
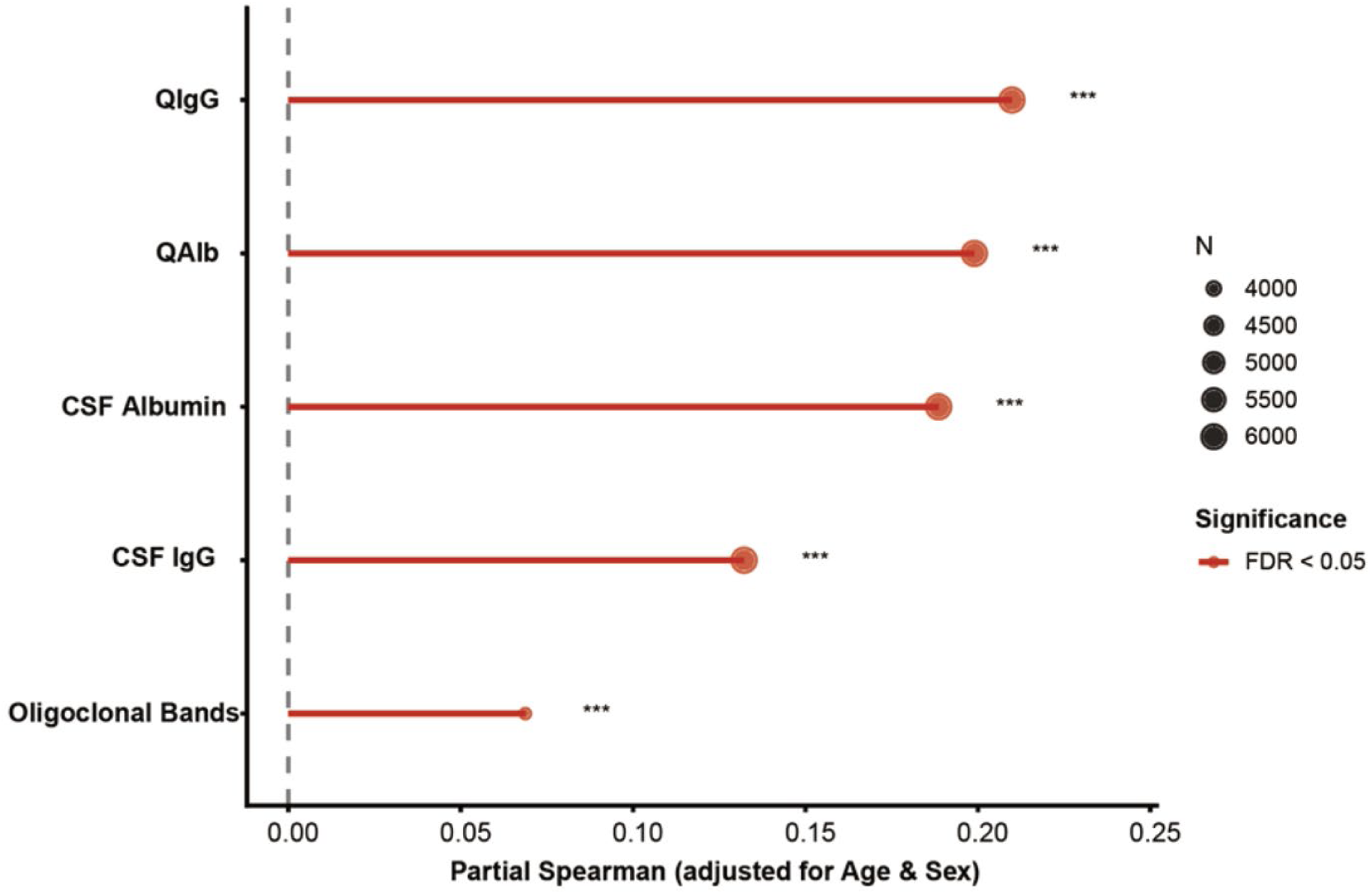
Associations between CSF barrier and intrathecal immune markers and CSF AgeAccel. Forest plot showing age- and sex-adjusted partial Spearman correlation coefficients between CSF AgeAccel and markers of BBB permeability and intrathecal immune activity, including QIgG, QAlb, CSF albumin, CSF IgG, and oligoclonal bands. Points indicate partial correlation coefficients, and horizontal bars represent 95% confidence intervals. Positive values indicate higher marker levels associated with greater CSF brain-age acceleration. Statistical significance was assessed using partial Spearman correlation analysis with false-discovery-rate (FDR) correction.

**Supplementary Figure 5.**
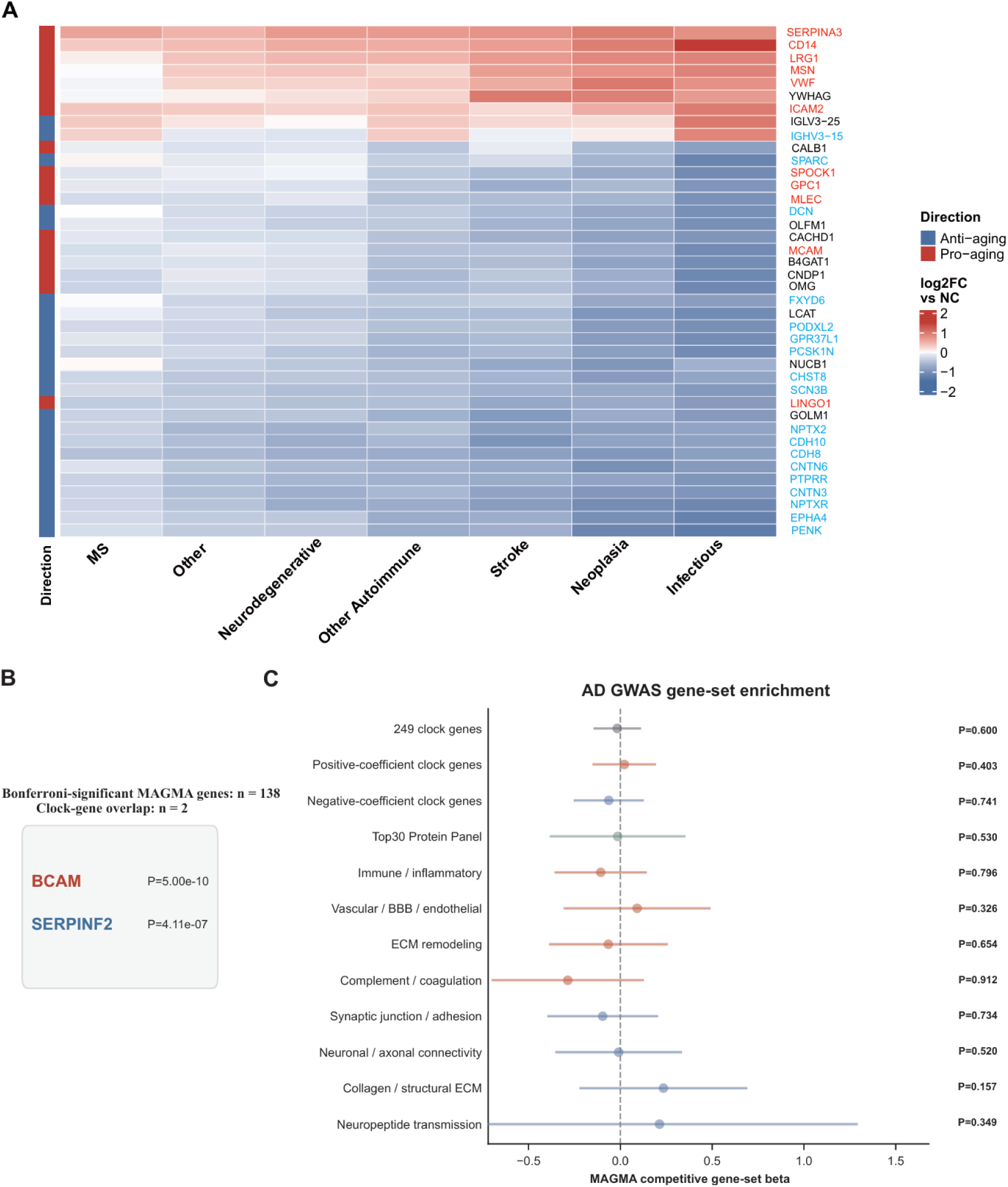
Disease-associated clock-protein dysregulation and limited Alzheimer’s disease GWAS enrichment. **(A)** Heatmap showing differential abundance patterns of selected CSF aging-clock proteins across neurological disease groups. For each disease group, protein abundance was compared with neurologically normal controls (NC). Displayed proteins were selected because they showed recurrent or prominent disease-associated changes across one or more disease categories. Colors indicate log2 fold change relative to NC, with red indicating higher abundance in the disease group and blue indicating lower abundance. The left side annotation denotes clock direction, with red indicating pro-aging proteins and blue indicating anti-aging proteins. Marked proteins overlap with the pathway-defined core proteins identified from the pro-aging and anti-aging module analyses, confirming that the disease-associated differential patterns are driven by the same two biological programs. **(B)** Direct overlap between CSF aging-clock genes and Bonferroni-significant MAGMA genes in the Alzheimer’s disease GWAS. Two clock genes overlapped with AD-associated MAGMA genes: BCAM and SERPINF2. **(C)** Competitive MAGMA gene-set enrichment of the same CSF aging-clock gene sets in the Alzheimer’s disease GWAS. No clock-derived gene set showed nominal enrichment.

**Supplementary Figure 6.**
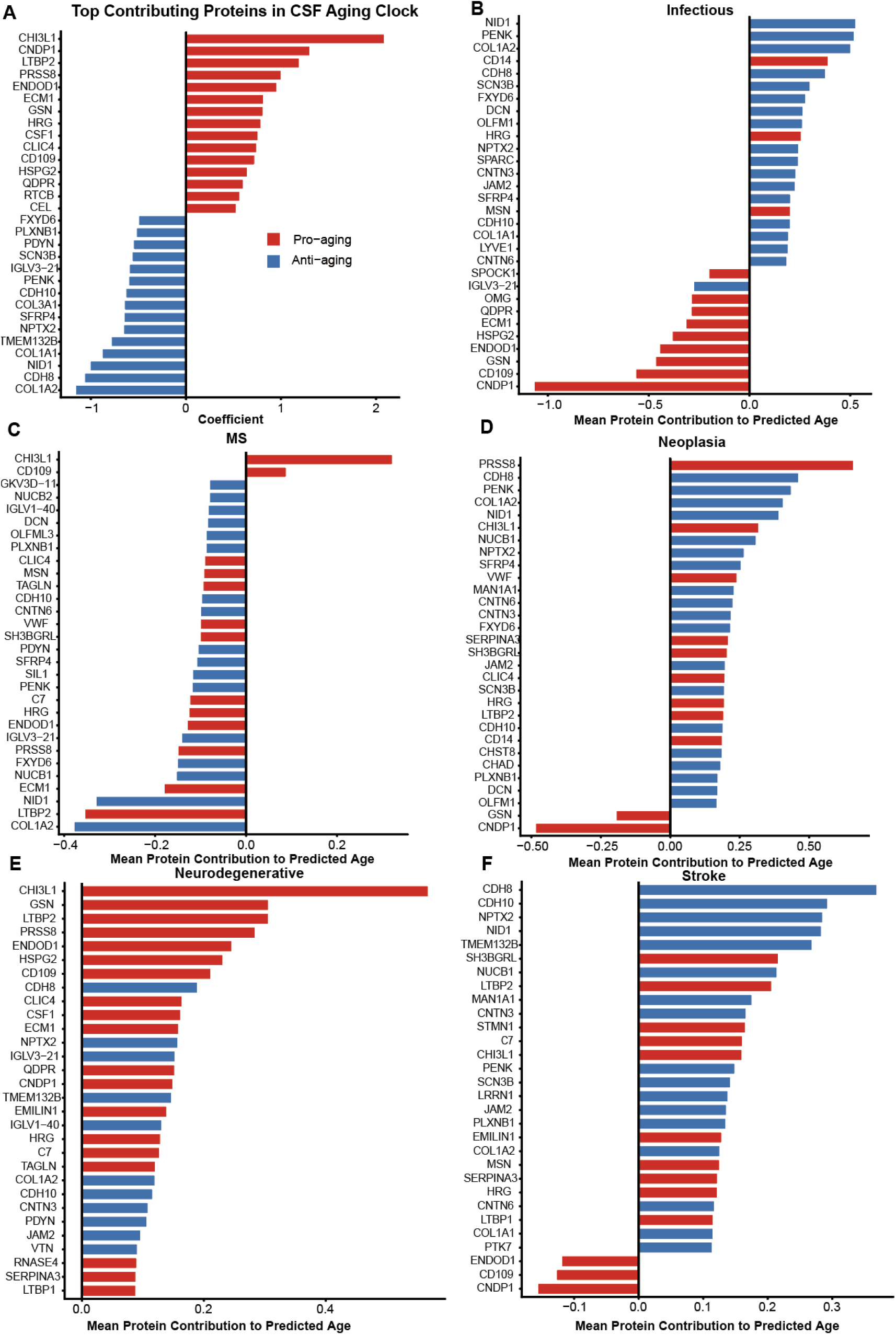
SHAP-based analysis of disease-specific protein contributions to the CSF proteomic aging clock. **(A)** Top 30 proteins contributing to the CSF proteomic aging clock, ranked by the absolute magnitude of elastic-net coefficients. Proteins with positive clock coefficients (pro-aging proteins) are shown in red, whereas proteins with negative clock coefficients (anti-aging proteins) are shown in blue. **(B–F)**Disease-specific SHAP (SHapley Additive exPlanations) contribution profiles in infectious diseases (B), multiple sclerosis (C), neoplasia (D), neurodegenerative diseases (E), and stroke (F). Bars represent mean SHAP values for individual clock proteins within each disease category. Positive SHAP values indicate contributions that increase predicted CSF proteomic age, whereas negative SHAP values indicate contributions that decrease predicted CSF proteomic age. Protein colors correspond to clock directionality, with red indicating pro-aging proteins and blue indicating anti-aging proteins. Labelled proteins denote the highest-contributing proteins within each disease category.

**Supplementary Figure 7.**
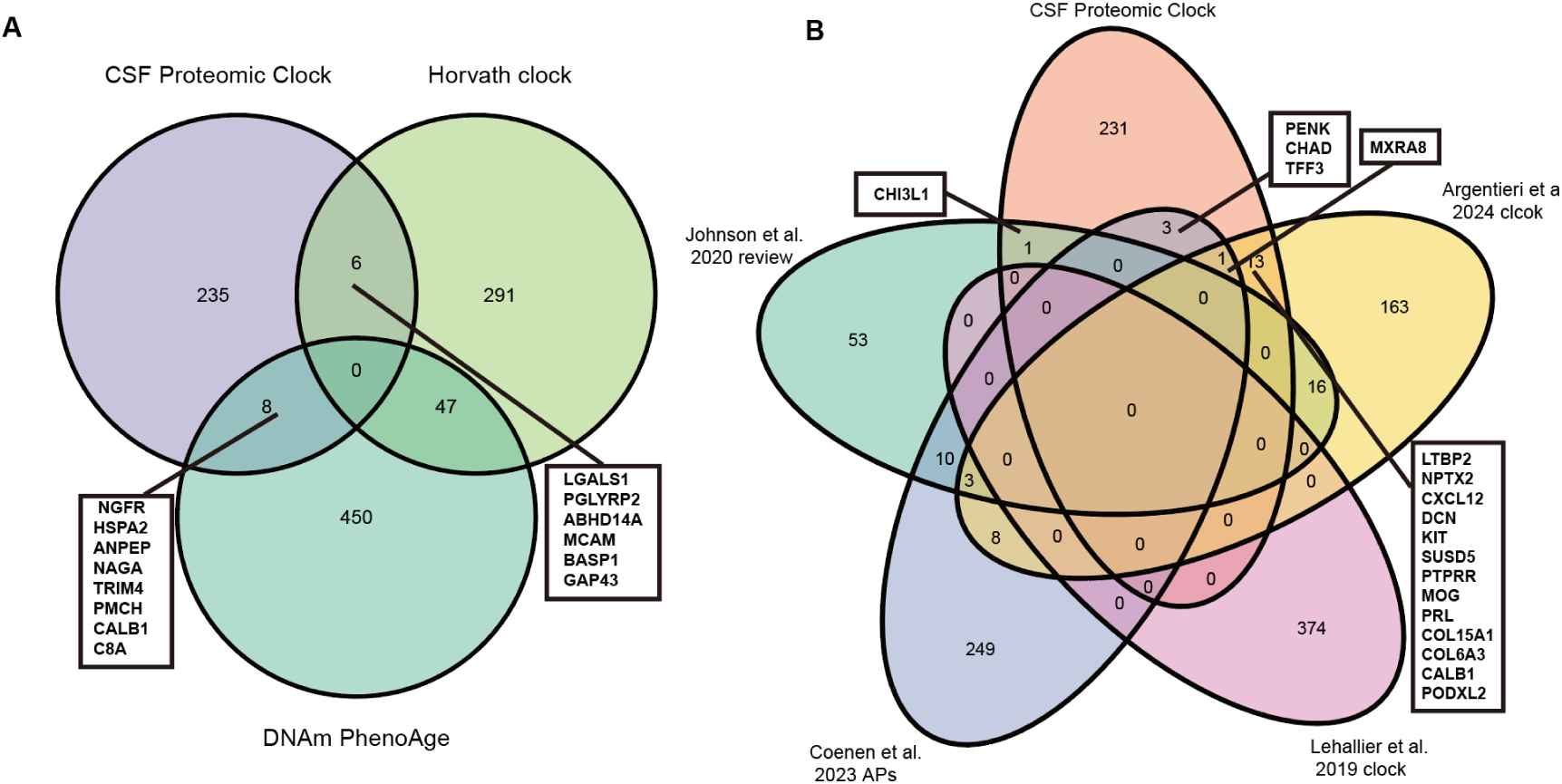
Limited overlap between the CSF proteomic aging clock and blood-based aging signatures. **(A)** Overlap between CSF clock proteins and genes represented in blood DNA methylation aging clocks (Horvath clock and DNAm PhenoAge)^13,14^. Numbers indicate unique and shared protein/gene counts. **(B)** Overlap between CSF clock proteins and proteins reported in published plasma proteomic aging clocks and age-associated protein signatures (Johnson et al., Coenen et al., Lehallier et al., Argentieri et al.)^7,15–17^. The majority of CSF clock proteins are not represented in any compared plasma proteomic aging dataset, indicating a largely CNS-specific molecular aging program.

**Supplementary Figure 8.**
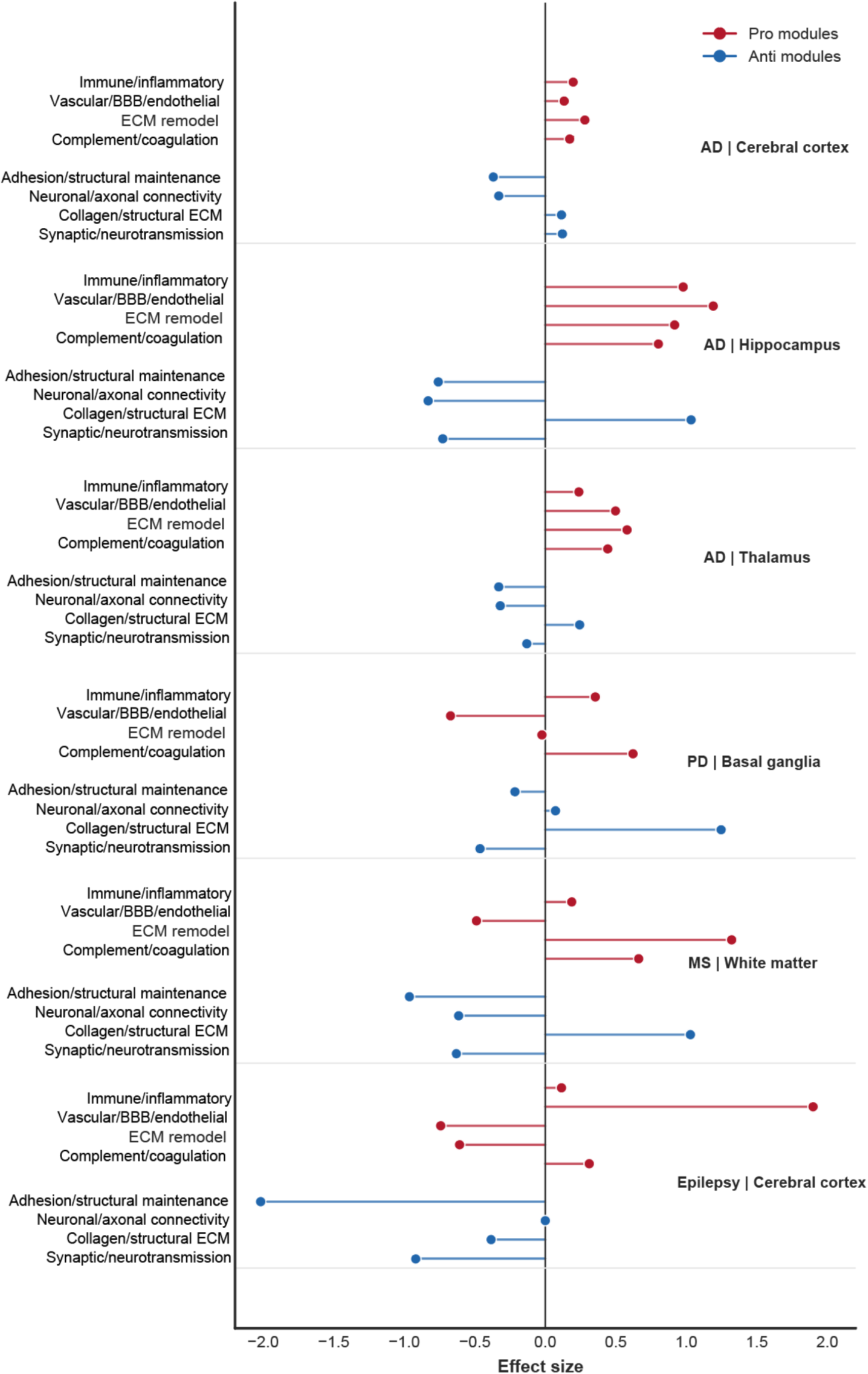
Module-level decomposition of disease-associated pro-aging and anti-aging programs across vulnerable brain regions. Effect sizes are shown as Cohen’s d comparing disease samples with region-matched healthy controls for each module score. Red indicates pro-aging modules, including immune/inflammatory, vascular/BBB/endothelial, ECM-remodeling and complement/coagulation programs. Blue indicates anti-aging modules, including adhesion/structural maintenance, neuronal/axonal connectivity, collagen/structural ECM and synaptic/neurotransmission programs. Positive values indicate higher module scores in disease samples relative to region-matched healthy controls, whereas negative values indicate lower scores. The collagen/structural ECM module showed a pattern distinct from neuronal anti-aging modules, with increased scores in several disease-relevant regions, supporting its evaluation as a separate remodeling component.

**Supplementary Figure 9.**
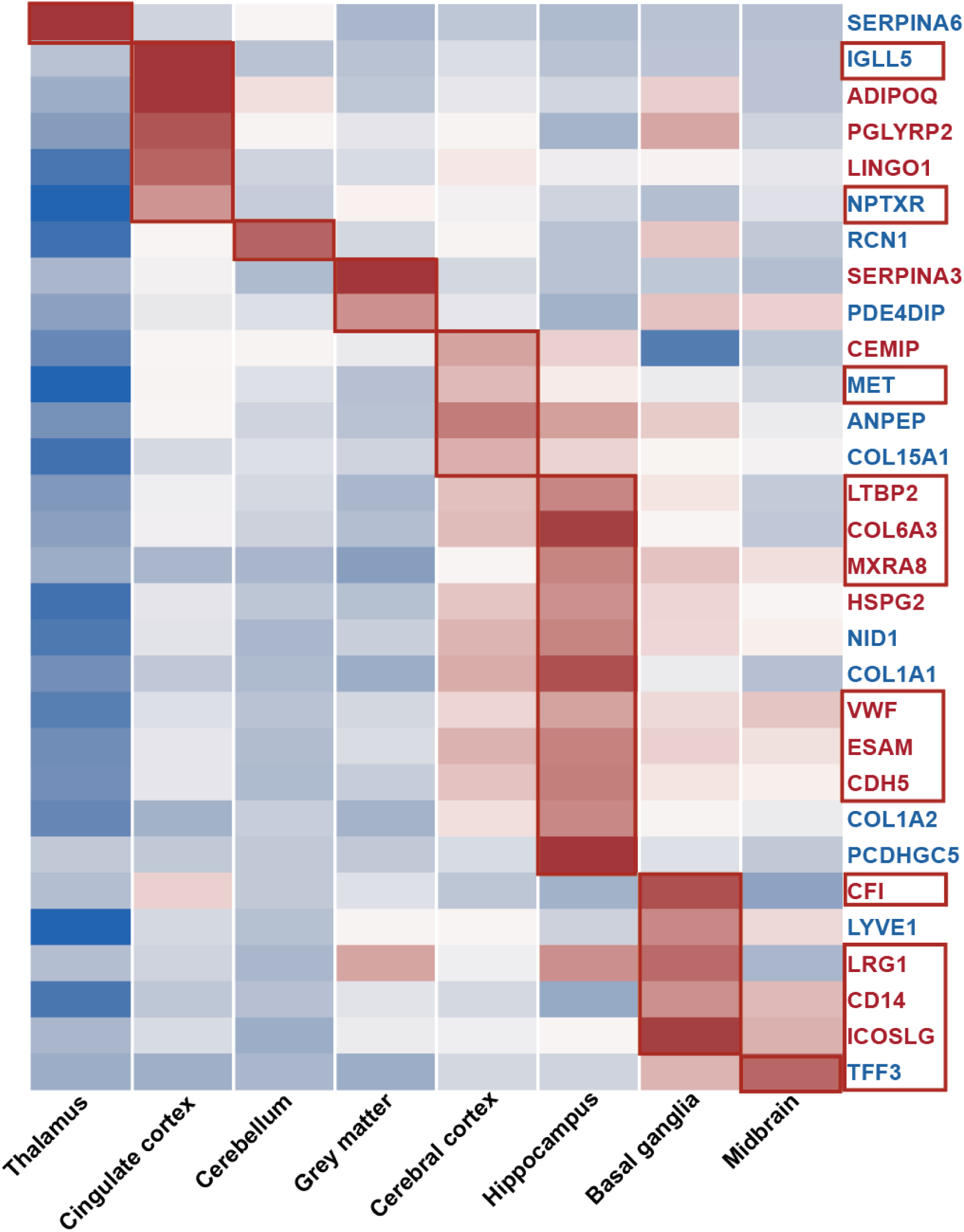
Region-specific expression heatmap of CSF aging-clock proteins in healthy human brain pseudobulk transcriptomic data. Rows represent clock proteins showing significant regional upregulation, defined by edgeR FDR < 0.05, logFC > 0.5, peak z-score ≥ 1 and peak z − second-highest z-score ≥ 0.3. Columns represent brain regions, and colors indicate row-scaled z-scores of log-CPM expression. Pro-aging proteins are labeled in red, and anti-aging proteins are labeled in blue. Genes are ordered according to their peak expression region. Boxes highlight proteins specifically enriched in each brain region. Proteins additionally showing age-associated expression in healthy brain samples with Pearson |r| > 0.3 and direction concordant with the CSF aging-clock direction were defined as healthy brain-region-specific aging benchmark proteins.

**Supplementary Figure 10.**
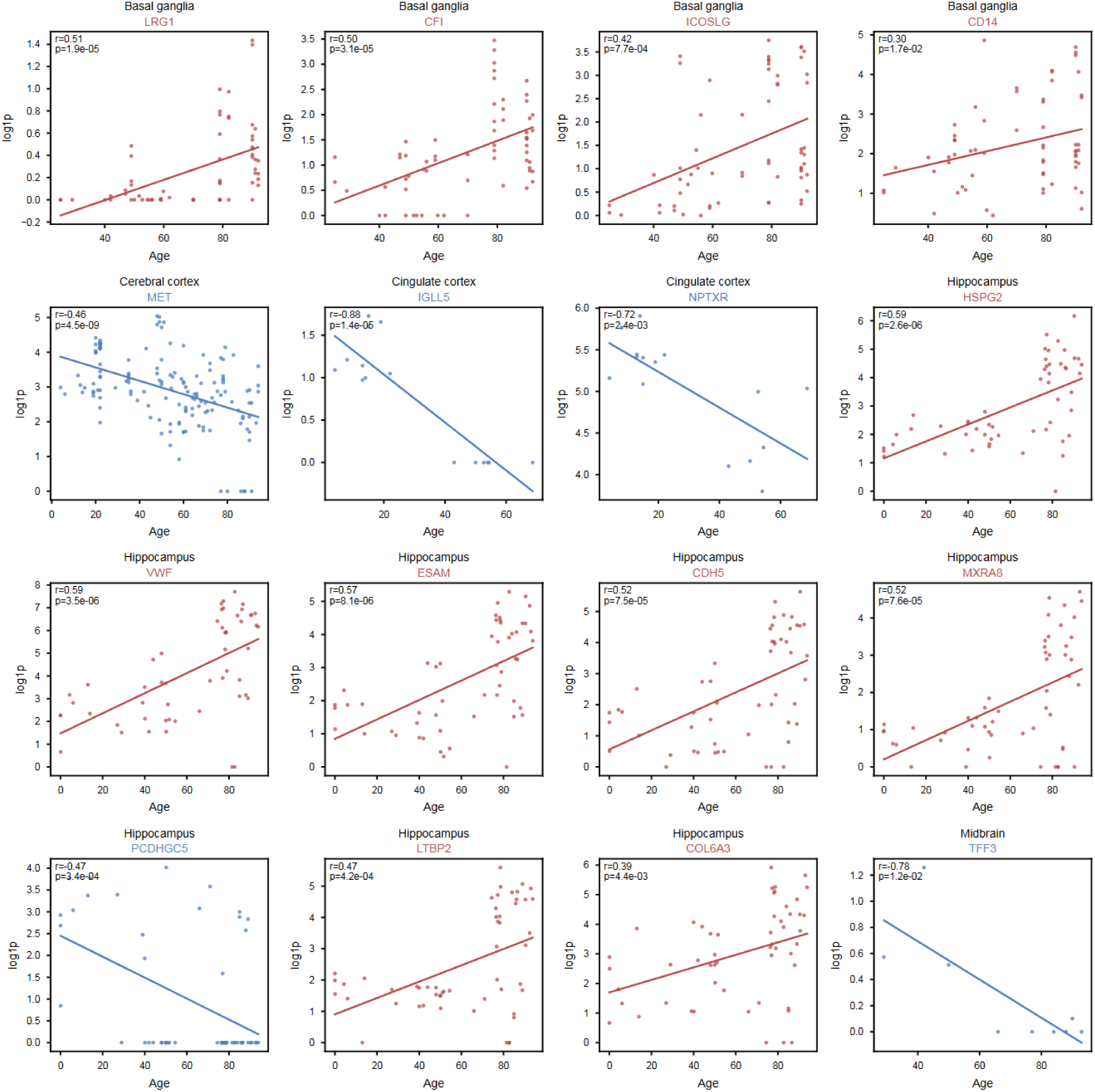
Age-associated expression of region-specific CSF aging-clock genes in the healthy human brain. Age information from a healthy human brain single-cell transcriptomic atlas was integrated with brain-region-level pseudobulk expression profiles. Pearson correlations between chronological age and gene expression were calculated for brain-region-specific CSF aging-clock genes within their corresponding anatomical regions. Genes showing significant age associations, directional concordance with the CSF aging-clock model, and absolute correlation coefficients greater than 0.3 were selected for visualization. Representative genes are shown for the basal ganglia, cerebral cortex, cingulate cortex, hippocampus, and midbrain. Red panels denote pro-aging genes with positive age-associated expression changes, whereas blue panels denote anti-aging genes with negative age-associated expression changes. Each point represents one healthy donor sample. Solid lines indicate linear regression fits. Pearson correlation coefficients (r) and corresponding P values are shown in each panel.

**Supplementary Figure 11.**
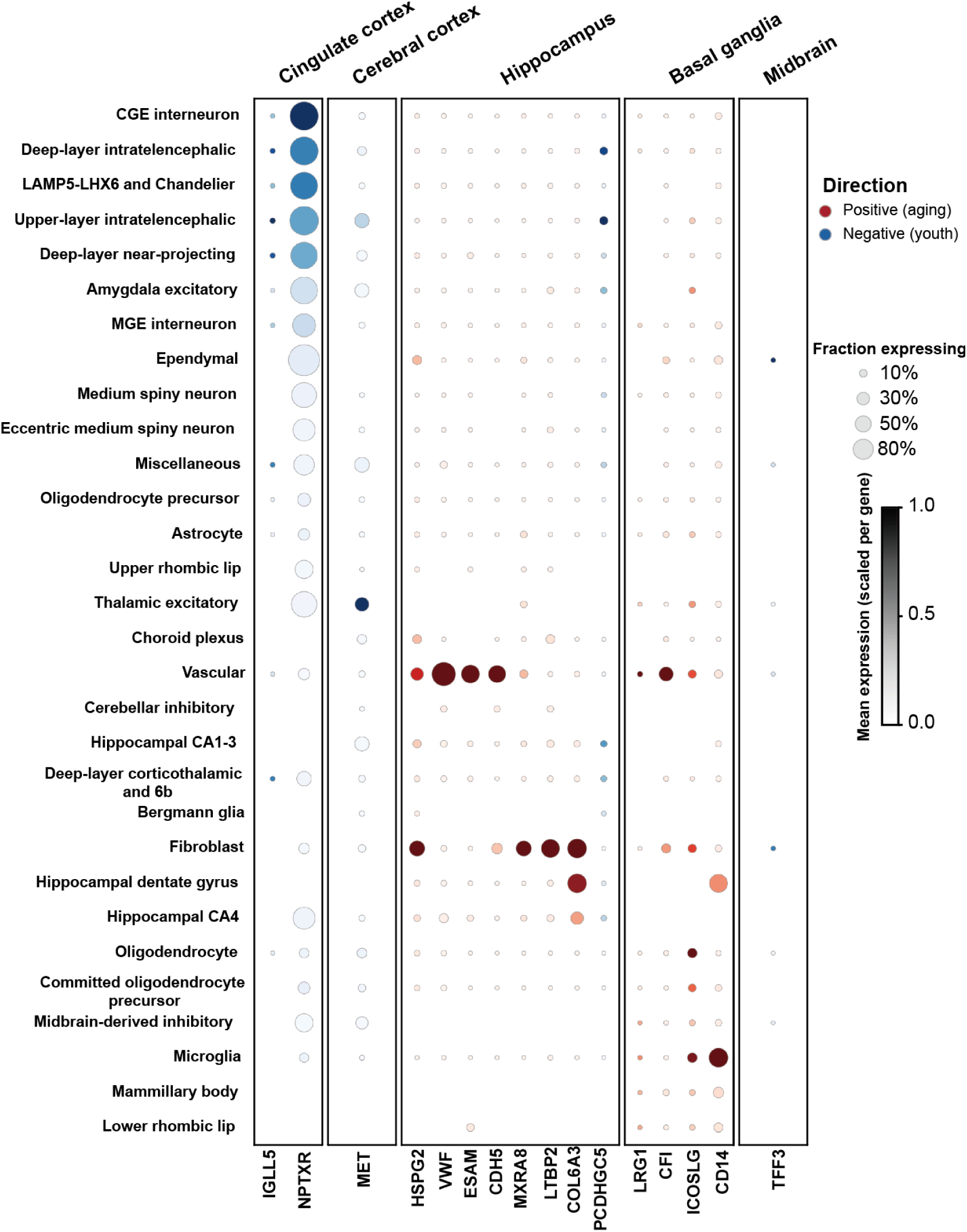
Cellular localization of the healthy brain-region-specific aging benchmark proteins identified. Dot plots show expression of these benchmark proteins across major brain cell populations and anatomical regions. Dot size represents the fraction of cells expressing each gene, and color intensity indicates scaled mean expression. This analysis links region-specific aging benchmark proteins to their likely cellular sources, including neuronal, glial, vascular and barrier-associated cell populations.

**Supplementary Figure 12.**
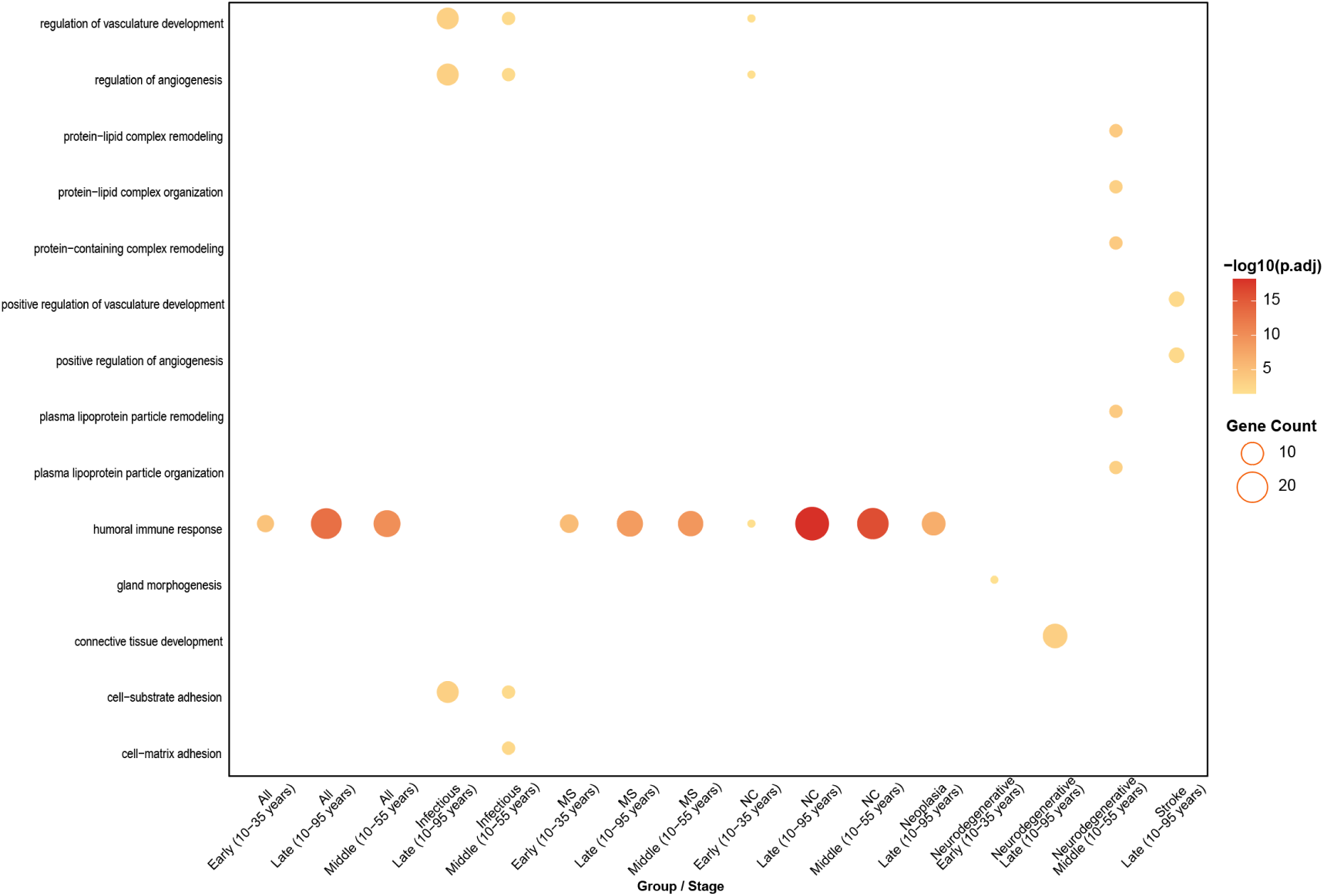
Biological pathways enriched among proteins upregulated across inflection-point-defined aging stages in healthy controls. Dot plot showing GO biological process enrichment among proteins that increased across aging stages defined by age-related inflection points in healthy-control CSF samples. The x-axis indicates pairwise comparisons between adjacent aging stages, and the y-axis shows significantly enriched biological processes. Dot color represents enrichment significance (−log10 adjusted P value), and dot size indicates the number of proteins contributing to each pathway. Enriched pathways included vascular and angiogenesis-related processes, such as regulation of vasculature development and regulation of angiogenesis, as well as immune-, extracellular matrix-, connective tissue-, and lipid-remodeling-related processes, including humoral immune response, connective tissue development, protein–lipid complex remodeling, and plasma lipoprotein particle remodeling.

**Supplementary Figure 13.**
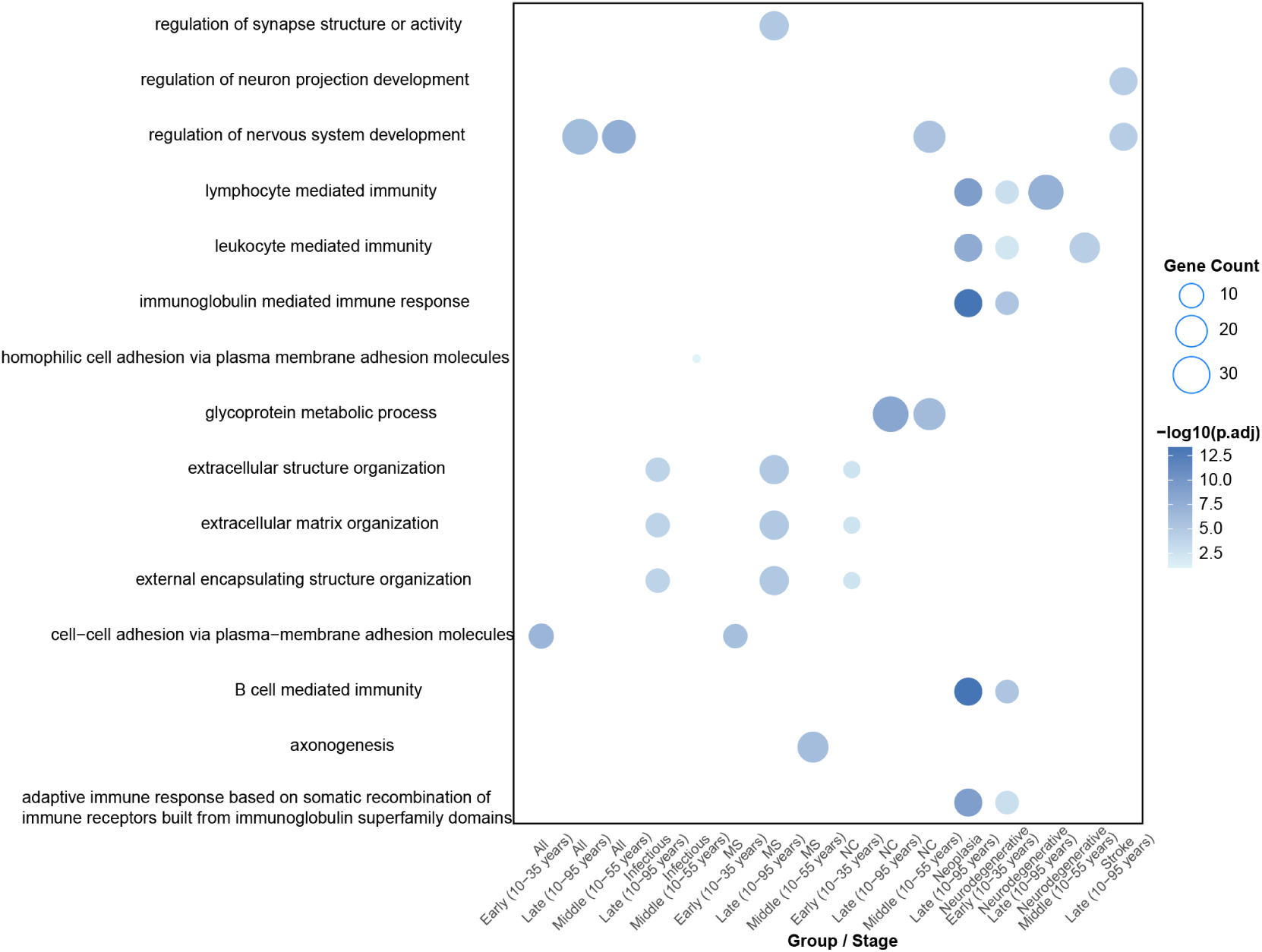
Biological pathways enriched among proteins downregulated across inflection-point-defined aging stages in healthy controls. Dot plot showing GO biological process enrichment among proteins that decreased across aging stages defined by age-related inflection points in healthy-control CSF samples. The x-axis indicates pairwise comparisons between adjacent aging stages, and the y-axis shows significantly enriched biological processes. Dot color represents enrichment significance (−log10 adjusted P value), and dot size indicates the number of proteins contributing to each pathway. Enriched pathways among downregulated proteins included neuronal and synaptic processes, such as regulation of synapse structure or activity, regulation of neuron projection development, regulation of nervous system development, and axonogenesis. Additional pathways were related to cell adhesion and extracellular structural organization, including homophilic cell adhesion via plasma membrane adhesion molecules, cell-cell adhesion, extracellular matrix organization, and extracellular structure organization. Immune-related pathways, including lymphocyte-mediated immunity, leukocyte-mediated immunity, B cell-mediated immunity, and immunoglobulin-mediated immune responses, were also represented in selected stage transitions.

